# Statistical learning dynamically shapes auditory perception

**DOI:** 10.1101/2024.09.09.612146

**Authors:** Sahil Luthra, Austin Luor, Adam T. Tierney, Frederic Dick, Lori L. Holt

## Abstract

Humans and other animals use information about how likely it is for something to happen. The absolute and relative probability of an event influences a remarkable breadth of behaviors, from foraging for food to comprehending linguistic constructions -- even when these probabilities are learned implicitly. It is less clear how, and under what circumstances, statistical learning of simple probabilities might drive changes in perception and cognition. Here, across a series of 29 experiments, we probe listeners’ sensitivity to task-irrelevant changes in the probability distribution of tones’ acoustic frequency across tone-in-noise detection and tone duration decisions. We observe that the task-irrelevant frequency distribution influences the ability to detect a sound and the speed with which perceptual decisions about its duration are made. The shape of the probability distribution, its range, and a tone’s relative position within that range impact observed patterns of suppression and enhancement of tone detection and decision making. Perceptual decisions are also modulated by a newly discovered perceptual bias, with lower frequencies in the distribution more often and more rapidly perceived as longer, and higher frequencies as shorter. Perception is sensitive to rapid distribution changes, but distributional learning from previous probability distributions also carries over. In fact, massed exposure to a single point along the dimension results in seemingly maladaptive loss of sensitivity - occurring entirely in the absence of feedback or reward – along a range of subsequently encountered frequencies. This points to a gain mechanism that suppresses sensitivity to regions along a perceptual dimension that are less likely to be encountered.

**Significance Statement:** Organisms as diverse as honeybees and humans pick up on probabilities in the world around them. People implicitly learn the likelihood of a color, price range, or even syntactic structure. How does statistical learning affect how we detect events and make decisions, especially when probabilities are completely irrelevant to the task at hand, and can change without warning? We find that people learn and track changes in perceptual probabilities irrelevant to a task and that this learning drives dynamic shifts in perception characterized by graded effects of enhancement – and primarily – suppression across acoustic frequency. This can result in a remarkably long-lived diminishment of perceptual sensitivity that seems maladaptive but may instead reflect use of likelihood to guide and sharpen perception.

## Introduction

We implicitly pick up information about the probability of white versus red cars on the road, the spatial position of objects in a room, and how likely different sounds might be within a soundscape – for instance, hearing a cow moo in a barnyard versus a hospital. The detailed distributional structure of sensory input leads us to expect some events and to be surprised by others.

How does distributional learning influence perception? Many studies have focused on learning across probabilistic input, whereby organisms implicitly discover regularities across continuous input dimensions (e.g., Love, 2003; McMurray, Aslin, Toscano, 2009; Rosenthal, Fusi, & Hochstein, 2001). For example, unsupervised learning of clustering in speech sounds may scaffold infants’ language acquisition (Werker, Yeung, & Yoshida, 2012; Cristià, 2011; Schatz et al., 2022). Other studies have focused on the outcome of distributional learning, manipulating stimulus probability to operationalize expectation and characterizing the influence of expectation on perception and neural representation (Summerfield & de Lange, 2014; Summerfield & Egner, 2009).

Some theoretical accounts of these expectation-driven effects predict that perception will prioritize high-probability expected input (consistent with Bayesian inference; de Lange et al., 2018). Indeed, frequent, expected stimuli are better detected than rare stimuli (Pinto et al., 2015; Stein & Peelen, 2015) and perceptual decisions about expected stimuli are speedier and more accurate (Summerfield & de Lange, 2014; Summerfield & Egner, 2009). This enhanced perception might be achieved via adjustments of weights on sensory channels that modulate gain, sharpening representation of frequent relative to rare input. Alternately, perceptual enhancements might be mediated by expectation-congruent memory representations (Summerfield & de Lange, 2014; Kok et al., 2012). Neuroimaging studies have revealed that representation of expected stimuli is enhanced via suppressed activity in voxels tuned away from expected stimuli (Kok et al., 2012; Yon et al., 2018).

Other accounts conclude, instead, that distributional learning accentuates infrequent, unexpected events (see Press et al., 2020). This prioritization is accomplished by suppressing expected input (Blakemore et al., 1998; Kilteni & Ehrsson, 2017; Richeter et al., 2018; Meyer & Olson, 2011; Kumar et al., 2017), leading to improved detection of rare stimuli (Milne et al., 2024). A third account posits that expectation can lead to enhancement in some contexts and suppression in others, with initial perceptual biases that tilt toward expected stimuli but can be cancelled out by highly surprising input (Press et al., 2020). But complicating matters, probability distributions experienced across a perceptual dimension may influence the bottom-up salience (Alink & Blank, 2021; Zivony & Eimer, 2024) or task relevance (Rungratsameetaweemana & Serences, 2019) of a dimension, each with the potential to impact perception. In sum, there is no consensus about how likelihood influences perception.

We propose that opposing theoretical perspectives may persist, at least in part, as a byproduct of empirical focus on dichotomous frequent-versus-rare likelihoods that necessarily limit the resolution with which the relationship between expectation and gain can be estimated. More complex probability distributions sampled across a continuous perceptual dimension have the potential to reveal granular, graded influences of expectation built from distributional learning across probability.

Here, we shape expectation by sampling stimuli probabilistically across the primary representational axis of the auditory system, acoustic frequency. Crucially, *acoustic frequency is task-irrelevant* across our 29 studies. This decouples expectation from task utility, unveiling the influence of distributional statistical learning across a task-irrelevant dimension on perception. We test how this learning impacts perception across unimodal, bimodal, and equiprobable distributions varying in statistical volatility and sampling density. Given evidence that distinct task goals can influence the impact of short-term input regularities on perception (Fritsche, Mostert, & de Lange, 2017), we use two tasks inspired by classic psychoacoustics literatures. One task examines detection of near-threshold tones in continuous noise. In an influential study, Greenberg and Larkin (1968) led listeners to expect a single constant-frequency tone to appear in noise but tone frequency varied on a minority of trials. Detection accuracy was superior for the expected, high-probability frequency with graded diminishment of detection accuracy as a function of distance from the expected frequency. This graded sensitivity has been interpreted as a frequency-selective attentional filter centered at the distribution mode (Scharf et al., 1987). Here, detection accuracy of near-threshold tones in noise provides a graded metric of the perceptual gain function arising from expectation built from distributional learning that supports directional assessment of enhancement versus suppression.

Complementing this, the speed of tone duration decisions reveals the influence of distributional learning across task-irrelevant acoustic frequency. Capitalizing on classic studies of Schröger and Wolf (1998) developed as a model of auditory distraction, participants decide whether a sound is “long” or “short” across tones with distinct and equiprobable durations. The tones’ acoustic frequency is task-irrelevant but carries a distributional regularity that impacts responses, with slower duration decisions consistent with longer processing time for tones with low-probability frequencies. Our parallel tests of tone-in-noise detection and tone duration decisions allow us to examine the influence of distributional learning on perceptual tasks possessing distinct processing demands.

To foreshadow, our distributional learning approach reveals influences of expectation that would be unobservable across simple frequency-versus-rare likelihoods. We find that distributional learning is not simply frequentist accumulation of events: equally likely events are differentially perceived as a function of their position within a probability distribution and are influenced by filter properties of both the sensory system and selective attention. We observe exquisite sensitivity to distribution shifts and robust carryover of influence from previously experienced distributions at multiple timescales. The influence of distributional learning on perception arises rapidly after a shift in distributions, but our data also provide evidence for ’stubborn predictions’ that linger (Yon, de Lange, & Press, 2019). In fact, massed exposure to a single point along the frequency dimension results in sustained diminishment of perceptual sensitivity of subsequently encountered frequencies as a function of their distance from that single point. This would seem maladaptive for perceptual precision but instead may reflect a mechanism that centers attentional gain around the most likely region(s) along a perceptual dimension. Across studies, the impact of distributional learning on perception is most consistent with a *sharpening* of representations through *suppression* of processing units tuned to less-likely stimuli (e.g., Yon et al., 2018).

## Results

Given the large number of experiments and results we report only test type, exact *p* values, and power for each statistical test in the main text; all tests are corrected for multiple comparisons. **Table S3** provides details on each reported analysis, including the relevant filename of the subject-wise data and analysis files available at https://osf.io/xdgnw/.

### Distributional learning alters the detection of tones in noise

We first ask whether distributional learning across a continuous sensory dimension affects the most basic perceptual process: detection. Does the probability with which a sound occurs influence the ability to hear it in noise?

For each detection study, individual detection thresholds are established immediately before the experiment using three iterations of a standard staircase technique adapted for online testing, Zhao et al., 2022; see **Materials and Methods**, **Fig 1a**, top). Thereafter listeners detect a tone presented at threshold in continuous white noise within one of two intervals (**Fig 1a**, bottom).

**Figure 1.**
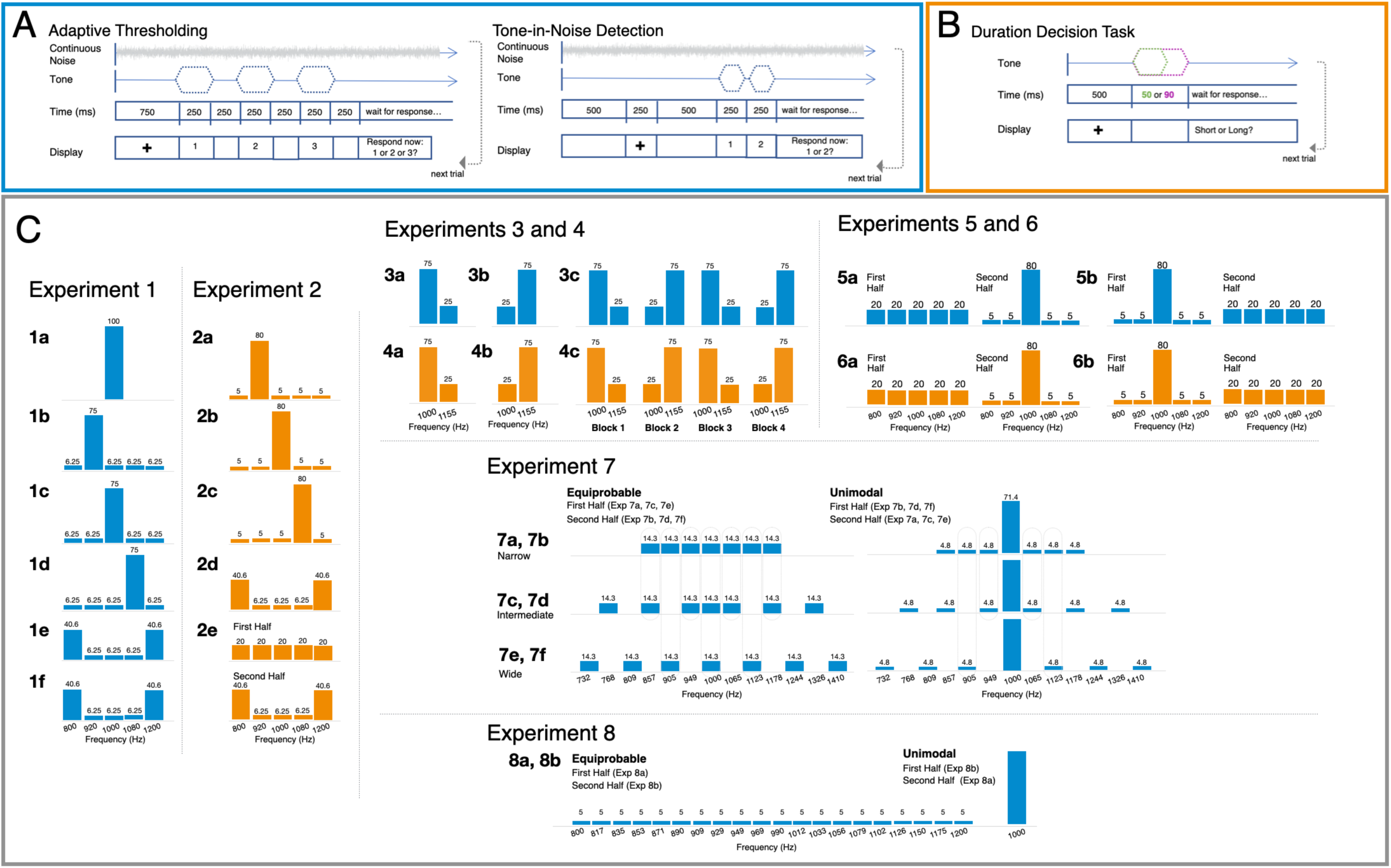
Tasks and Distributional Regularities. **(A)** The tone-in-noise detection task involved two phases: adaptive threshold estimation followed by the tone-in-noise detection task. Threshold estimation trials began with continuous noise and a fixation cross (750 ms), after which a 1000-Hz tone was presented with equal probability in one of three 250-ms detection windows (250 ms ISI), each indicated by a number (1, 2, or 3) on the screen. A prompt 250-ms after the third detection window elicited participants’ report of the interval containing a tone (here, shown in the first interval). Tone intensity followed the 3-down, 1-up procedure to estimate 79% accuracy (see **Methods and Materials**). The noise continued through the tone-in-noise detection task, shown in the bottom of **(A)**. For each trial, 500 ms preceded a 250 ms fixation cross and another 500 ms period. A 250-ms sinewave tone with intensity + 0.75 dB above the threshold estimated in the adaptive thresholding task appeared in one of two 250-ms intervals (250 ms ISI), indicated by a “1” and a “2" on the screen, respectively. Participants reported which interval contained the tone (here, shown as interval 2). Tone frequency varied according to the distributions in **(C)**. **(B)** In the duration-decision task, each trial involved a 1000-ms fixation followed by a 50 or 90 ms sine wave tone (equal probability) and participants reported “long” or “short” with a button press. **(C)** Probability distributions for each experiment, as a function of acoustic frequency. Blue distributions indicate tone-in-noise detection experiments. Orange distributions indicate duration-decision experiments.

Exp 1a establishes baseline detection accuracy when a single acoustic frequency (1000 Hz) is 100% probable. Exp 1b-f draw from a pool of five easily differentiable frequencies (800, 920, 1000, 1080, 1200 Hz) spaced ∼13x the just-noticeable difference in frequency (Sek & Moore, 1995). In Exp 1b-d, one highly probable frequency comprises 75% of the 320 trials. The remaining four tones each occur on just 6.25% of trials, creating a unimodal distribution across frequency. Exp 1e has a bimodal probability distribution with 800 Hz and 1200 Hz frequencies each presented on 40.6% of trials, with each other frequency presented on 6.25% of trials. Exp 1f is identical to Exp 1e, except that the frequency for threshold estimation is 1080 Hz, rather than 1000 Hz as in Exp 1a-e. **Fig 1c** illustrates these distributions across the acoustic frequency dimension.

In Exp 1, stimulus probability strongly modulates tone detection in noise across Exp 1b-f with better detection of high-probability frequencies at the distribution mode (**Fig 2a**; ANOVA, Freq x Exp interaction, p = 1.761x10^-31^, η^2^ = 0.169). Detection of only 1000 Hz (Exp 1a: 100% probability; average accuracy 77.9%) does not differ from detection of the highest-probability frequency in unimodal distributions (Exp 1b-d: 75% probability; average accuracy 75.3%; ANOVA, p = 0.242, η^2^ = 0.012). But detection of the modal frequencies in the bimodal distributions (40% probable) is lower than when a single frequency is 80% or 100% probable (Exp 1e-f: 40.6% probability; average accuracy 70.3%; ANOVA, p = 0.006, η^2^ = 0.053 versus Exp 1b-d, ANOVA, p = 0.003, η^2^

**Figure 2.**
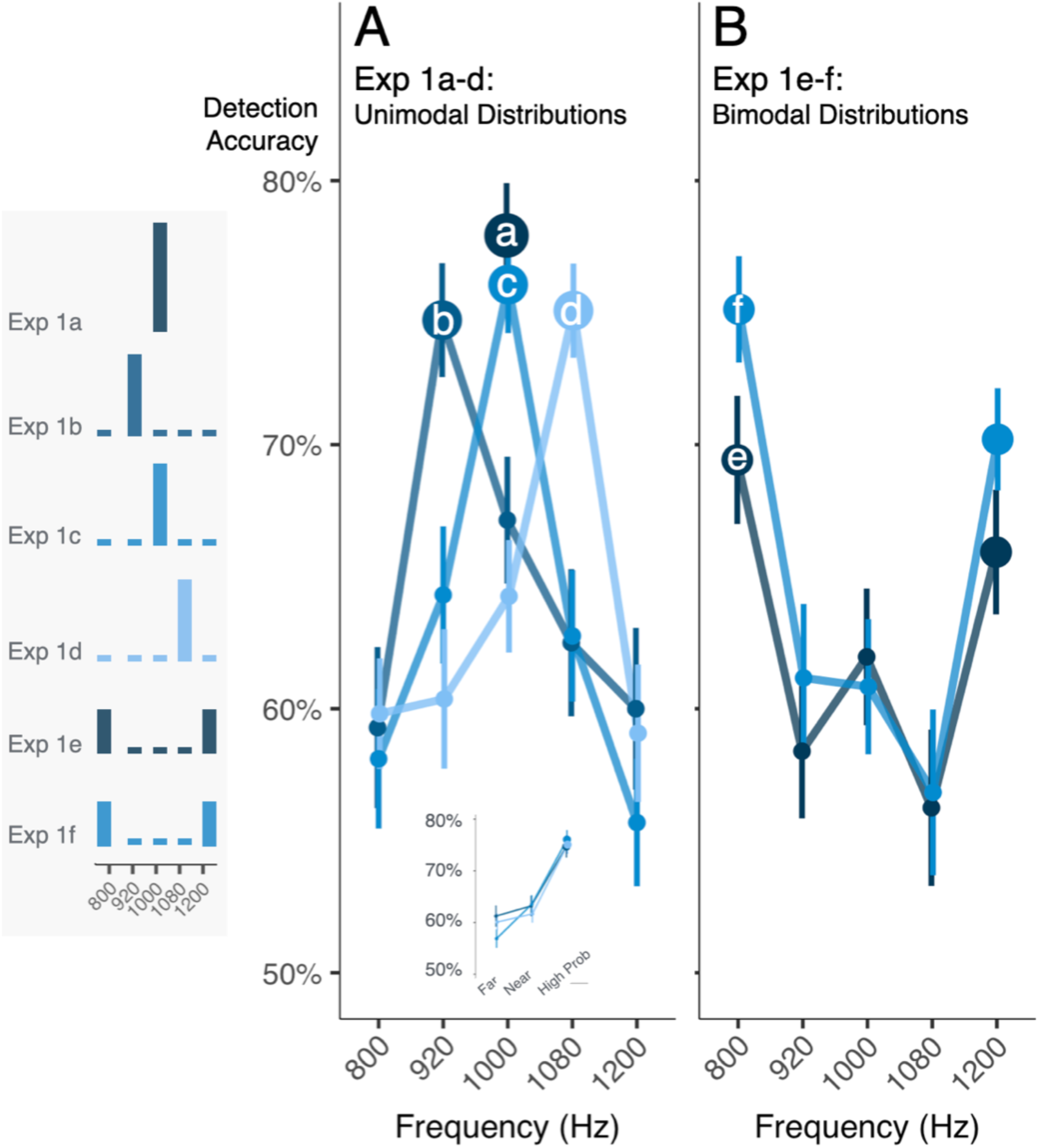
Distributional learning alters the detection of tones in noise. Each panel plots mean detection accuracy as a function of tones’ acoustic frequency. The histograms to the left show distributional regularities for each experiment. Marker size scales with tone probability. Error bars are standard error of the mean. **(A)** Detection accuracy for a single-point distribution at 1000 Hz in Exp 1a approximates the expected detection accuracy estimated by the preceding threshold procedures and serves as a reference baseline for single frequency detection. For Exp 1b-d the distribution mode is detected best, with equivalently low-probability tones detected more poorly as a function of distance from the mode (see inset). **(B)** Bimodal distributions produce a ‘dual spotlight’ with detection accuracy best at the modes. Exp 1e-f differ only in the frequency used to estimate the threshold (1000 and 1080 Hz, respectively).

= 0.101 versus Exp 1a).

Proximity to the high probability tone also influences detection (**Fig 2a**). The low-probability frequencies of Exp 1b-d share the same probability, yet those closer to a high-probability frequency are better detected than those further away (ANOVA, p = 0.014, η^2^ = 0.022). When the high-probability frequency is centered in the range of frequencies defining the distribution, this graded detection accuracy difference is symmetric (near > far to high-probability frequency, ANOVA, p = 0.004, η^2^ = 0.259). When the high-probability frequency is nearer to the distribution edge (Exp 1b and Exp 1d), there is an asymmetric detection curve (ANOVA, p = 0.015, η^2^ = 0.034): a sharp detection decrement toward the distribution edge is contrasted with a more gradual ‘ski slope’ decrement toward the middle of the frequency range (see inset, **Fig 2a**). In sum, equiprobable rare tones are detected more accurately if they are adjacent to the distribution mode, but this advantage is modulated by the position of the high probability tone relative to the range of the frequency distribution.

These results must be understood in the context of the operating characteristics of the auditory system, for which there is a critical bandwidth of approximately 1/6 of an octave within which the cochlea has limited ability to resolve stimuli (Fletcher, 1940). At 1000 Hz, the critical bandwidth is ∼130 Hz. Thus, taking Exp 1c (1000 Hz mode) as an example, these sensory filter properties likely contribute to the ‘rescue’ of detection of low-probability tones with frequencies situated close to the distribution mode (920, 1080 Hz). Even so, the asymmetry of the shallow ‘ski slope’ decrement toward the middle of the frequency range compared to the steep decrement toward the edges of the range in Exp 1b and Exp 1d suggests that other mechanisms are also at play. Where the high probability tone is situated within the range of experienced frequencies also has an influence. If effects were driven solely by integration within an auditory critical band, we should not observe such contextual influence.

More complex probability distributions also modulate detection (**Fig 2b**). Exp 1e shows that a bimodal probability distribution with higher-probability (40.6%) frequencies at the edges of the distribution (800 and 1200 Hz) induces a ‘dual spotlight’ across the frequency dimension. Listeners detect the higher-probability tones more accurately than neighboring low-probability tones (920 and 1080 Hz, linear contrast, Bonferroni-corrected p = 3.590 x 10^-5^, Cohen’s d = 0.757) with a marginal difference compared to the middle 1000 Hz tone (linear contrast as above, p = 0.072, Cohen’s d = 0.418).

Note that for Exp 1e, detection of 1000 Hz tones has a numerical (but not significant) detection advantage compared to the other low-probability tones (**Fig 2b**). Two ‘spotlights’ centered at the high-probability tone frequencies would yield a “V” rather than this observed “W” detection profile. We speculated that the numerical detection advantage for 1000 Hz might arise from experience with 1000 Hz in the 90-trial threshold-setting procedure that precedes Exp 1e. Exp 1f falsifies this hypothesis. Changing the initial threshold-setting frequency to 1080 Hz elicits a similar “W” profile and, importantly, replicates the overall ’dual spotlight’ at 800 and 1200 Hz (linear contrast as above, p = 4.62 x 10^-8^, Cohen’s d = 0.985, **Fig 2b**).

In summary, Exp 1 demonstrates that distributional learning modulates sound detection. Replicating and extending classic studies in psychoacoustics (Greenberg & Larkin, 1968), tones with higher-probability frequencies are better detected in noise than lower-probability frequencies. The impact of distributional learning is graded across frequency, with better detection of low-probability frequencies that lie closer to high-probability frequencies than equally improbable, but more distant, frequencies. This effect is further influenced by the overall distributional context: the protective effect of proximity to the high-probability tone depends on its position within the range of encountered frequencies. Moreover, bimodal distributions with two higher-probability frequencies at the edges of the frequency range elicit a ‘dual spotlight’.

### Distributional learning across a task-irrelevant dimension impacts perceptual decisions

Listeners track probabilities across acoustic frequency despite the irrelevance of frequency to the Exp 1 detection task. Previous findings show that similar probability distribution manipulations affect perceptual decision response times (Schröger & Wolff, 1998; Garrido, Dolan & Sahani, 2011). We next ask whether statistical learning over a probability distribution defined across task-irrelevant *frequency* impacts the time course of decisions about a sound’s *duration*.

In Exp 2a-c, participants report whether a tone is long or short, with 50 ms and 90 ms tones presented equiprobably across 400 trials (**Fig 1b**; see **Materials and Methods**). Task-irrelevant tone frequency varies across five frequencies (800-1200 Hz) in the manner of Exp 1 (**Fig 1c**). There are four improbable tone frequencies (each 5% of trials), and a single probable frequency (80% of 400 trials, Exp 2a: 920 Hz; Exp 2b, 1000 Hz; Exp 2c: 1080 Hz). In Exp 2d, 800 Hz and 1200 Hz are presented on 40.625% of trials with the other frequencies each presented on 6.25% of trials to create a bimodal distribution (320 trials). In Exp 2e, the five tones are equiprobable (20%) across the first half of the study and then switch to the bimodal distribution of Exp 2d (640 total trials).

Across Exp 2a-c, the probability of a tone’s *frequency* significantly impacts the speed of *duration* decisions (ANOVA, p = 7.620 x 10^-7^, η^2^ = 0.017, **Fig 3a**). Response times (RTs) are slower for tones with low, compared to high, probability frequencies (linear contrast, unpooled error, p =1.628 x 10^-17^, Cohen’s d = 0.516). Further, RTs for duration decisions to equiprobably rare frequencies are graded as a function of their distance from the high-probability distribution mode. Compared to RTs to the most probable frequency, those to the adjacent low-probability frequencies are slower (Holm-corrected post hoc comparison, p = 4.316 x 10^-12^, Cohen’s d = 0.387) and frequencies furthest away from the high-probability frequency are yet slower than the frequencies nearer the high-probability one (Holm-corrected p = 5.137 x 10^-6^, Cohen’s d = 0.258). (These patterns hold true for each Exp 2a-c study, p < .05 Holm-corrected). This replicates and extends classic observations from psychoacoustics (Schröger & Wolff, 1998) and mirrors the graded influence on Exp 1 detection accuracy (**Fig 2a**).

**Figure 3.**
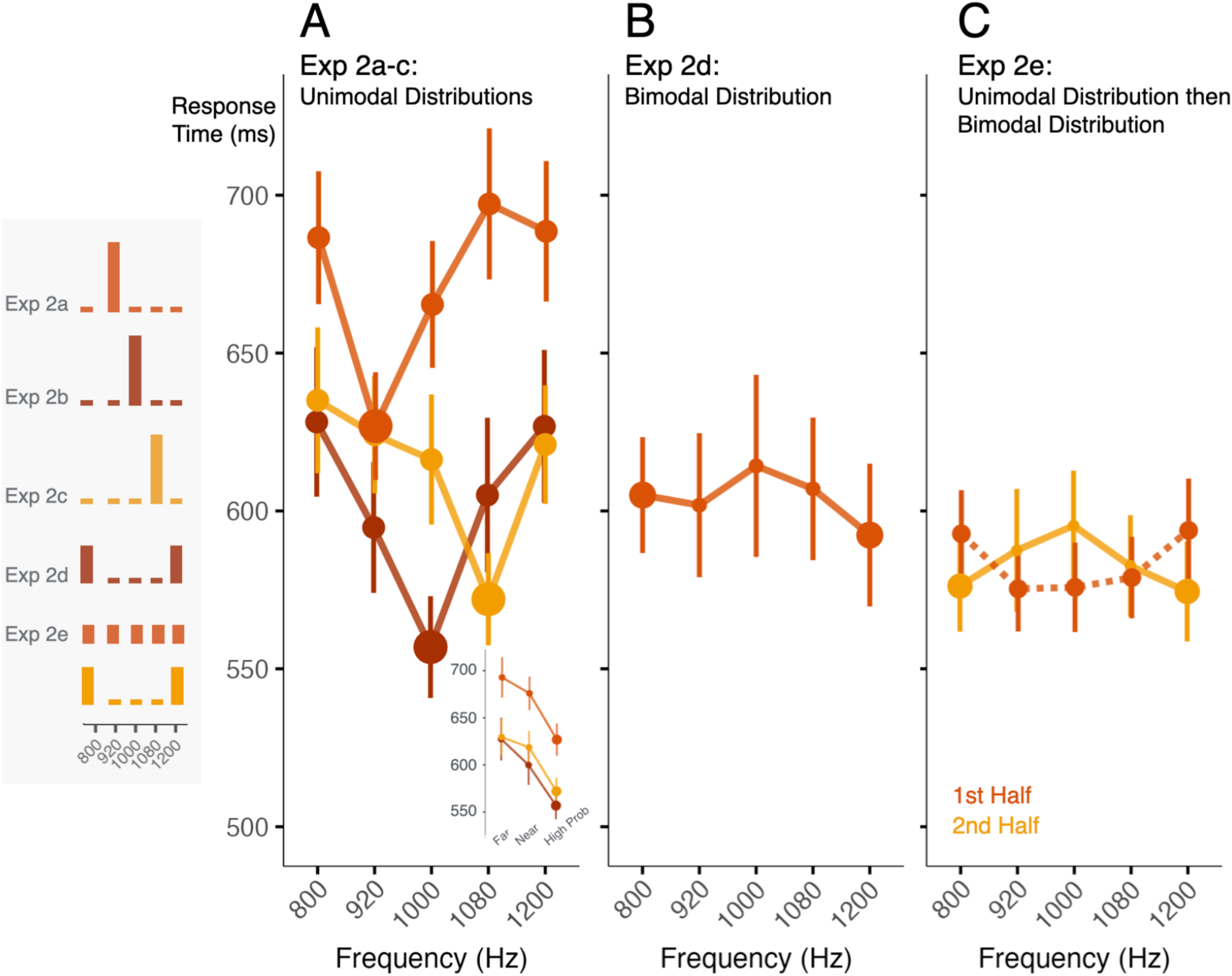
Distributional learning across a task-irrelevant dimension impacts perceptual decisions. Each panel plots mean response time as a function of tones’ acoustic frequency. The histograms to the left show distributional regularities for each experiment. Marker size scales with tone probability. Error bars are standard error of the mean. **(A)** Response time to report tone duration is impacted by the probability of tones’ acoustic frequency across Exp 2a-c. The influence is graded, with faster decision times for equivalently low-probability tones closer to the distribution mode (see inset). **(B)** Unlike the dual spotlight for tone detection in Exp 1e-f, there is no significant response time difference for the two more probable modes in Exp 2d, a consequence of a frequency-duration perceptual bias (see **Fig S1**). **(C)** Exp 2e evaluated the frequency-duration bias across an equiprobable distribution in the first half of the study (orange, dashed) with a switch to the bimodal distribution at study midpoint (yellow, solid). The bias is largest at the edges of the distribution where it interacts with the bimodal distributional regularity (see **Fig S1**).

However, unlike the dual spotlight for tone detection in Exp 1e-f, there is no significant RT advantage for making duration decisions about the higher-probability 800 and 1200 Hz tones in Exp 2d (**Fig 3b**; ANOVA effect of frequency, p = 0.526, η^2^ = 0.010). To examine this more closely, Exp 2e introduces a distribution change: five initially equiprobable (20%) frequencies (320 trials) shift to mirror the Exp 2d bimodal distribution mid-study (320 trials; see **Fig 1c**). This allows us to characterize potential frequency-duration interactions that may exist, independent of probability. Indeed, and unexpectedly, when tone frequencies are equiprobable in the first half of trials, decision RTs are *longer* for 800 Hz and 1200 Hz compared to other frequencies (ANOVA on Frequency, p = 0.031, η^2^ = 0.026; post-hoc linear contrast p = 0.002, Cohen’s d = 0.205) (**Fig 3c**).

Investigating this reveals a *novel frequency-duration perceptual bias*: duration decisions for lower-frequency tones (800, 920 Hz) are more accurate and faster for long (90 ms) compared to short (50 ms) tones whereas those for the highest frequency tone (1200 Hz) are more accurate and faster for short compared to long tones (**Fig S1**; ANOVA, Frequency x Duration interaction, RT: p = 0.003, η^2^ = 0.029; Accuracy (Acc): p = 3.738 x 10^-5^, η^2^ = 0.037). This perceptual bias is mirrored, but only qualitatively, in Exp 2d (**Fig S1**; p > 0.05, η^2^ = 0.010, with lower frequencies related to longer durations and higher frequencies with shorter durations). Notably, the bias is largest at the edges of the frequency distribution (800 and 1200 Hz) where it interacts with the bimodal distribution modes of Exp 2d-e, helping to explain why the dual spotlight observed in Exp 1e-f detection is not apparent in Exp 2d duration decisions. When we inspect the data from Exp 2a-c (**Fig S1**) we also observe the longer-lower/shorter-higher bias in the context of the unimodal distributions (ANOVA, Frequency x Duration interaction, RT: p = 3.968 x 10^-6^, η^2^ = 0.020; Acc: p = 0.003, η^2^ = 0.020). In other words, listeners found it easier to identify long durations when tones were relatively lower in frequency; conversely, it was easier to identify short durations when the sound was a relatively higher frequency tone. This impacted response time and interacted with the probability manipulation.

In summary, distributional learning across a task-irrelevant dimension affects perceptual decisions. The speed with which participants report the *duration* of a tone is impacted by the *probability of the tone’s frequency*. As with tone detection in noise in Exp 1a-f, learning across the probability distribution produces a graded influence on perceptual decisions: decisions across equivalently low-probability tones differ as a function of the tone’s distance in frequency from a high-probability tone. Moreover, Exp 2 demonstrates that seemingly intrinsic biases across acoustic dimensions may influence and/or disguise the impact of short-term statistical input regularities (for other examples see Roark & Holt, 2022; Bröker et al., 2024). These “intrinsic” biases might arise from distributional learning across longer timescales or perhaps sensory processing (see **Discussion**), and interact with short-term distributional regularities as shown in Exp 2a-e.

### Perceptual sensitivity and decisions rapidly update in volatile statistical contexts

Studies of statistical learning often investigate static distributions. But real-world environments can be volatile: for example, listeners often encounter talkers speaking different accents with different distributional regularities. The perceptual weight of different speech cues can rapidly alter in response to shifts in distributional regularities (e.g., Hodson et al., 2023; Murphy et al., 2023). It is not clear whether fundamental processes like detection and perceptual decisions are modulated by statistical volatility across *task-irrelevant* sensory dimensions.

Here, across six studies, we examine distributions composed of two tones: one high probability frequency and one low probability frequency (**Fig 1c**), akin to dichotomous probability distributions often used in studies of expectation and attention (e.g., Zivony & Eimer, 2024). In Exp 3a-b (detection) and Exp 4a-b (duration decision) we examine static two-frequency distributions to assure that effects of distributional learning observed across 5-tone distributions in Exp 1 and Exp 2 hold even in the simplest 2-tone sensory context over 320 trials. Exp 3a and Exp 4a examine detection and duration decisions, respectively, with 1000 Hz occurring across 75% of trials and 1155 Hz occurring over the remaining 25% of trials. Exp 3b and Exp 4b examine detection and duration across the complementary probability distribution. In Exp 3c and Exp 4c, we model a dynamic statistical context where these two-frequency distributions alternate every 160 trials. Participants experience four 160-trial blocks, with 1000 Hz high-probability (75%) and 1155 Hz low-probability (25%) in the first block, and probabilities alternating across frequencies in subsequent blocks.

Across Exp 3a and Exp 3b, we find equal and opposite effects of frequency probability, with the high probability tone detected on average ∼6% more accurately than the low probability tone (**Fig 4a**; ANOVA, Freq x Prob interaction, p = 3.361 x 10^-6^, η^2^ = 0.073). In Exp 4a and Exp 4b, RTs to the high probability tone frequency are on average ∼28 ms faster than those to the low-probability frequency (**Fig 4b**, ANOVA, p = 1.375 x 10^-6^, η^2^ = 0.010). We also observe the perceptual ’low-frequency à long-duration / high frequency à short-duration’ bias of Exp 2 even in this dichotomous probability distribution, with faster RTs for long-low/short-high duration-to-frequency pairings (ANOVA, Freq x Duration interaction, RT: p = 9.34 x 10^-6^, η^2^ = 0.013; Acc: p = 6.318 x 10^-5^, η^2^ = 0.023). In summary, a 2-tone frequency probability distribution affects tone in noise detection. It also affects individuals’ speed in making perceptual decisions across a different, task-relevant input dimension, but this effect is modulated by pre-existing perceptual biases.

**Figure 4.**
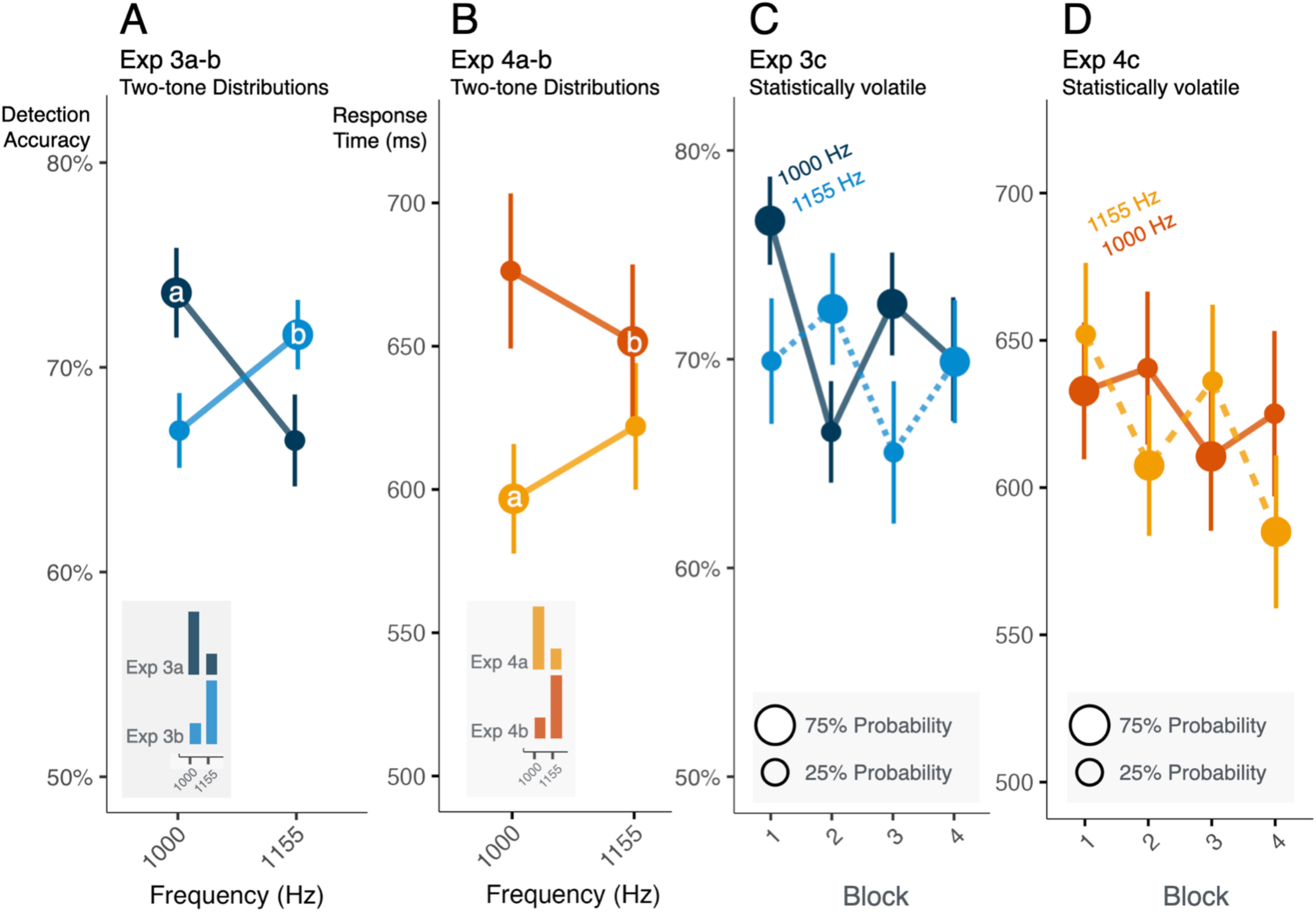
Perceptual sensitivity and decisions rapidly update in volatile statistical contexts. For Exp 3a-c mean detection accuracy as a function of acoustic frequency is plotted in blue; for Exp 4a-c mean response times in the duration-decision task are plotted in orange. Marker size scales with tone probability. In **(A)** and **(B)** the insets show the probability distributions. In **(C)** and **(D)** color indicates the tone frequency and marker size indicates its probability. Error bars are standard error of the mean. **(A)** Probability distributions defined across just two acoustic frequencies impact tone detection, with more accurate detection for high-probability tones in Exp 3a-b. **(B)** Two-tone distributions defined across task-irrelevant acoustic frequency also impact the response time to make duration decisions, with slower decisions to low-probability tones in Exp 4a-b. **(C)** As tone probability shifts every 160 trials across four blocks in Exp 3c, detection is more accurate for the high-probability, compared to low-probability, tones. **(D)** Similarly, in Exp 4c, the speed of duration decisions mirrors volatile probability changes: lower probability tone frequencies elicit slower decisions.

In the statistically volatile context established in Exp 3c, there is a detection advantage for the more probable frequency, with significant ‘flips’ in detection accuracy due to short-term reversals in tone probability for the first three blocks of Exp 3c (**Fig 4c**; ANOVA, Freq x Block interaction, p = 2.495 x 10^-5^, η^2^ = 0.099, each block p < 0.05). In the final block, there is no significant difference in detection accuracy across frequencies.

Likewise, transient changes in probability distribution affect the efficiency of perceptual decisions in Exp 4c (**Fig 4d**, ANOVA, Freq x Block interaction, p = 5.253 x 10^-7^, η^2^ = 0.040). RTs are slowest for the less probable frequency in all blocks (all p < 0.05 Bonferroni-corrected). Even in this dynamic context we again observe the systematic frequency-duration perceptual bias discovered in Exp 2 (ANOVA, Freq x Duration interaction, RT: p = 0.019, η^2^ = 0.039; Acc: p = 0.019, η^2^ = 0.077).

In summary, probability distributions defined across two acoustic frequencies elicit implicit statistical learning that impacts perception. The influence is rapid: probability exerts its influence across just 160 trials. As input statistics change, implicit statistical learning influences sound detection and perceptual decision making.

### The influence of distributional learning is consistent with a gain mechanism exhibiting hysteresis

We observe strong influences of distributional learning across unimodal probability distributions on detection accuracy and the speed of duration decisions (Exp 1 and Exp 2) that holds for dichotomous probabilities and follows volatile statistics across an experiment (Exp 3 and Exp 4). Here in Exp 5 (detection) and Exp 6 (duration decisions), we borrow from the distribution-switch design established in Exp 2e (**Fig 1c**). This distribution manipulation enables us to investigate how distributional learning influences detection and duration decisions across a changing statistical context. Moreover, by establishing perception across equiprobable distributions as a baseline, we reveal granular and graded changes in detection and decision making that emerge as distributional learning builds expectations, including enhancement and suppression of expected stimuli.

With equiprobable frequencies in the first half of Exp 5a, detection accuracy is consistent across frequency (**Fig 5a**; overall ∼65%, with unexpectedly better detection for 800 Hz, ANOVA, p = 0.009, η^2^ = 0.129). In the second half of Exp 5a, probabilities shift to mirror Exp 1b (1000 Hz 75%; all others 6.25%). This shift drives changes in accuracy which differ across frequencies (p = 8.511 x 10^-7^, η^2^ = 0.142). The 1000 Hz tones, which are now more probable, are better detected than they were in the first (equiprobable) half of Exp 5a (p = 0.002, Cohen’s d = 0.648), whereas the frequencies nearest (marginal p = 0.058, Cohen’s d = 0.554) and furthest (p = 0.027, Cohen’s d = 0.727) from 1000 Hz, which are now less probable, are more poorly detected than they were in the first half of the study (all Bonferroni-corrected linear contrasts with pooled error).

**Figure 5.**
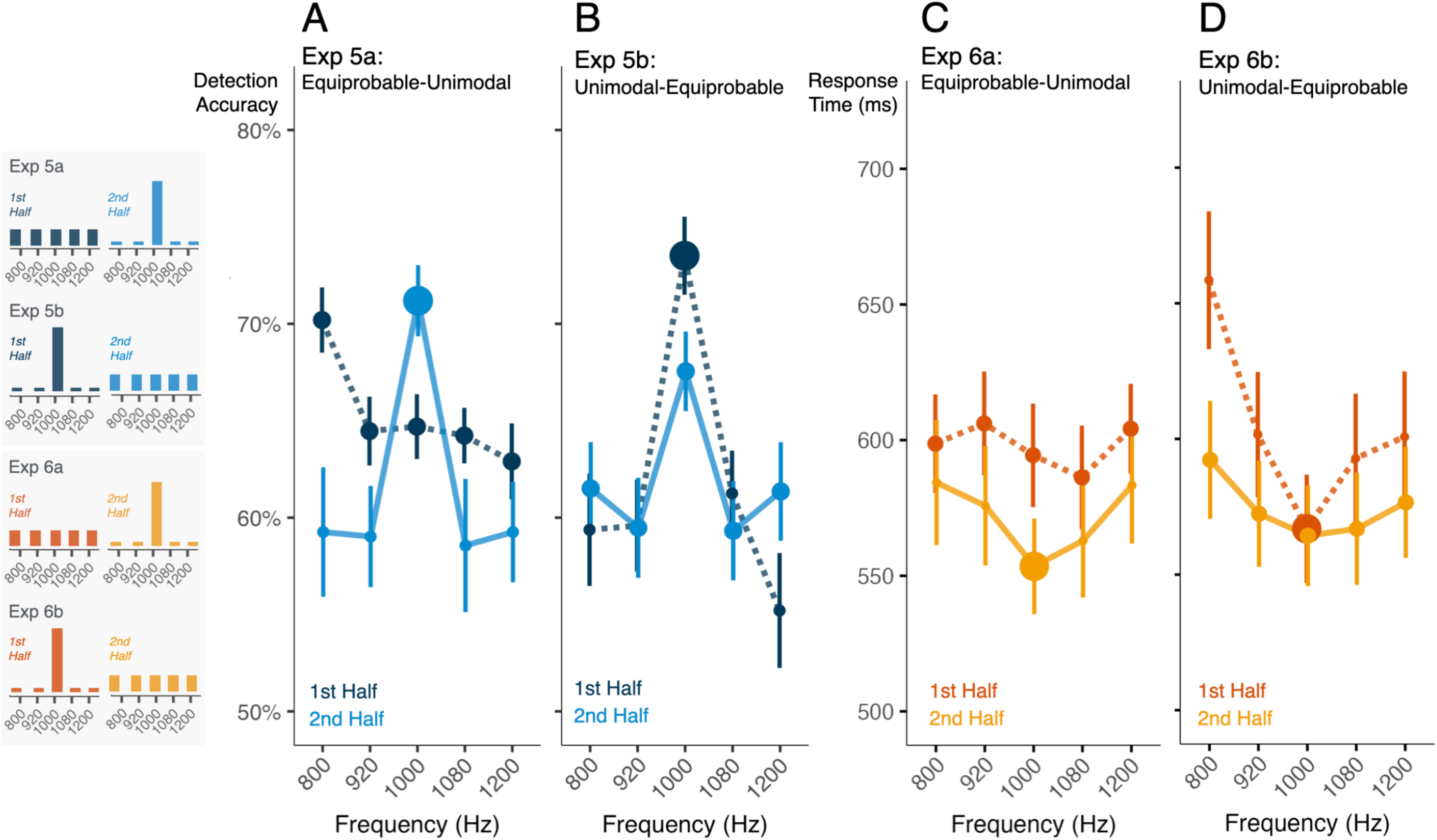
The influence of distributional learning is consistent with a gain mechanism exhibiting hysteresis. In Exp 5a-b mean detection accuracy as a function of acoustic frequency is plotted in blue; in Exp 6a-b mean response times in the duration-decision task are plotted in orange. The histograms to the left show distributional regularities for each experiment. Marker size scales with tone probability. In each panel, the darker color (dotted line) indicates behavior in the first half of the experiment; the lighter color (solid line) indicates behavior in the second half, when distributional regularities shift. Error bars are standard error of the mean. **(A)** Exp 5a establishes detection accuracy across a equiprobable distribution, then shifts to a unimodal distribution centered on 1000 Hz. Detection accuracy improves for the distribution mode with increased probability and decreases for frequencies with decreased probability. **(B)** Exp 5a switches from a unimodal distribution centered at 1000 Hz to an equiprobable distribution. Note the hysteresis at 1000 Hz, where detection remains elevated even into the second half of the study. **(C)** In Exp 6a, duration-decision times are flat with equiprobable frequencies in the first half. Introduction of a unimodal distribution centered at 1000 Hz leads to faster duration decisions at the mode. **(D)** In Exp 6b the unimodal distribution shifts to equiprobable at the study midpoint and response times shift substantially; note that this effect interacts with the frequency-duration bias identified in Exp 2.

In Exp 5b, we reverse distribution order. With a unimodal distribution centered on 1000 Hz in the first half of Exp 5b, detection generally resembles Exp 1c (**Fig 5b**), with better accuracy for high-probability 1000 Hz compared to low-probability frequencies (Bonferroni-corrected linear contrast, p = 1.023 x 10^-7^, Cohen’s d = 1.255), but with only a numerical detection advantage for frequencies nearest (920 and 1080 Hz) versus furthest (800 and 1200 Hz) from the probable center frequency (p = 0.262, Cohen’s d = 0.267, Bonferroni-corrected). When tone frequencies become equiprobable mid-study, again the probability shift drives differential changes in accuracy (p = 1.815 x 10^-4,^ η^2^ = 0.056). Here, the influence of the unimodal distribution carries over to confer a detection advantage to 1000 Hz, which was formerly highly probable, compared to other frequencies, which were formerly less probable (Bonferroni-corrected linear contrast, p = 1.383 x 10^-4^, Cohen’s d = 0.577). Detection of 1000 Hz tones decreased in accuracy from the first to the second study half due to the probability shift (p = 1.162 x 10^-5^, Cohen’s d = 0.496), but detection accuracy for the formerly low-probability tones did not change, despite a more than 3-fold probability increase (p = 1, Bonferroni corrected).

In sum, statistical learning across a unimodal distribution provokes a persistent effect on detection. For example, in Exp 5b, the initially highly probable 1000 Hz tone continued to be detected more accurately than other tones even after tone frequencies became equiprobable. Conversely, the tones adjacent 1000 Hz, which were initially relatively improbable, continued to be detected poorly even after the shift to the equiprobable distribution. Next, we use this distribution shift design to examine duration decisions.

Exp 6a begins with equiprobable frequencies and shifts mid-study to a unimodal distribution centered at 1000 Hz (80%, each other frequency 5%; **Fig 1c**). Exp 6b reverses this order. In the first half of Exp 6a, RTs in the duration-decision task across equiprobable frequencies are similar (**Fig 5c**, ANOVA, p = 0.163, η^2^ = 0.018). When probabilities shift to a unimodal distribution centered on 1000 Hz mid-study, RTs drop overall (ANOVA, p = 0.011, η^2^ = 0.061). Although there is a numerical ’V-shaped’ RT advantage for the now-probable 1000 Hz compared to increasingly more distant frequencies, this pattern does not differ significantly from the first half of the experiment (ANOVA, p = 0.245, η^2^ = 0.005).

In the first, unimodal probability half of Exp 6b, decision response times exhibit the “V” shape around the high-probability 1000 Hz tone also observed in Exp 2b (ANOVA, effect of frequency, p =6.847 x 10^-8^, η^2^ = 0.080, **Fig 5d**). Decisions about low-probability frequencies near to 1000 Hz are only numerically slowed compared to 1000 Hz itself (this and following test Bonferroni-corrected linear contrasts, p = 0.058, Cohen’s d = 0.227) but faster than to those further away from 1000 Hz (p = 0.014, Cohen’s d = 0.299).

When all frequencies become equiprobable mid-study in Exp 6b, there is a change in the degree to which frequency modulates decision RTs (ANOVA, p = 0.024, η^2^ = 0.017), but the 1000 Hz decision advantage persists in the second half (**Fig 5d**). Even though 1000 Hz is now 20% probable, RTs are not significantly different than in the first experiment half when it was 80% probable (linear contrast, p = 0.720, Cohen’s d = 0.025). Like detection in Exp 5b, there is carryover from experience with the unimodal distribution in the first half of the study, such that decision RTs are still modulated by frequency (ANOVA, p = 8.306 x 10^-5^, η^2^ = 0.068). RTs to report decisions for 1000 Hz continue to be significantly faster than for the now-equally-probable far frequencies (p = 0.020, Cohen’s d = 0.199), although not significantly faster than nearby frequencies (p = 0.668, Cohen’s d = 0.054). Finally, we again observe the duration-frequency bias established in the prior duration-decision studies (ANOVA, Freq x Duration interaction, RT: p = 1.608 x 10^-4,^ η^2^ = 0.013; Acc: p = 0.006, η^2^ = 0.021).

In summary, the impact of distributional learning on both detection and perceptual decisions emerges quickly and exhibits hysteresis, persisting even after the unimodal probability distribution flattens so that tones are equiprobable.

### The detailed shape of statistically-driven gain is modulated by range, distribution, and sampling density

In Exp 7, we make a more in-depth exploration of how expectations built up from distributional learning are impacted by statistical context, including frequency range and sampling density. Across six tone-in-noise detection studies, Exp 7 provides detailed information about the shape of the gain that emerges from distributional learning and how it evolves after an abrupt change in distributional statistics. We use these within-experiment distributional changes to estimate the emergence of enhancement and suppression of frequencies via distributional learning.

Exp 7a-f incorporate a mid-study change in distribution from equiprobable to unimodal or vice versa. The studies vary the range and density of 7 tone frequencies that define the distributions (**Fig 1c**) from ***narrow*** (Exp 7a,b; 5.5 semitone range), ***intermediate*** (Exp 7c,d; 9.47 semitones range), to ***wide*** (Exp 7e,f; 11.36 semitone range). In each range, frequencies are symmetrically arranged around 1000 Hz (like Exp 1c). As in prior studies, we group frequencies according to their distance (near, middle, and far) from the center frequency, which changes from highly probable to equiprobable or vice versa. In Exp 7a,c,e, the 7 frequencies are equiprobable (14.3%) until the experiment mid-point when 1000 Hz tones comprise the majority (71.4%) of trials and the other six tones are lower probability (4.8%). This order is reversed in Exp 7b,d,f. Below, we first describe detection accuracy patterns separately for Exp 7a,c,e (equiprobable to unimodal) and Exp 7b,d,f (unimodal to equiprobable), and then aggregate detection data across the unimodal conditions from each experiment to maximize power to detect effects of statistical context.

In Exp 7a,c,e, an equiprobable distribution precedes a switch to a unimodal distribution centered on 1000 Hz (see **Fig 6a-c**). Across these three studies, detection accuracy in the equiprobable first halves does not vary across frequency (ANOVA, p = 0.399, η^2^ = 0.003), nor is it modulated by the different frequency ranges across Exp 7a,c,e (ANOVA, p = 0.115, η^2^ = 0.035), and there is no interaction of frequency and range (p = 0.119, η^2^ = 0.011). Average detection accuracy across these equiprobable distributions is 64%, which does not differ significantly from that of the 5-frequency equiprobable distribution of Exp 5 (ANOVA, p = 0.219, η^2^ = 0.039).

**Figure 6.**
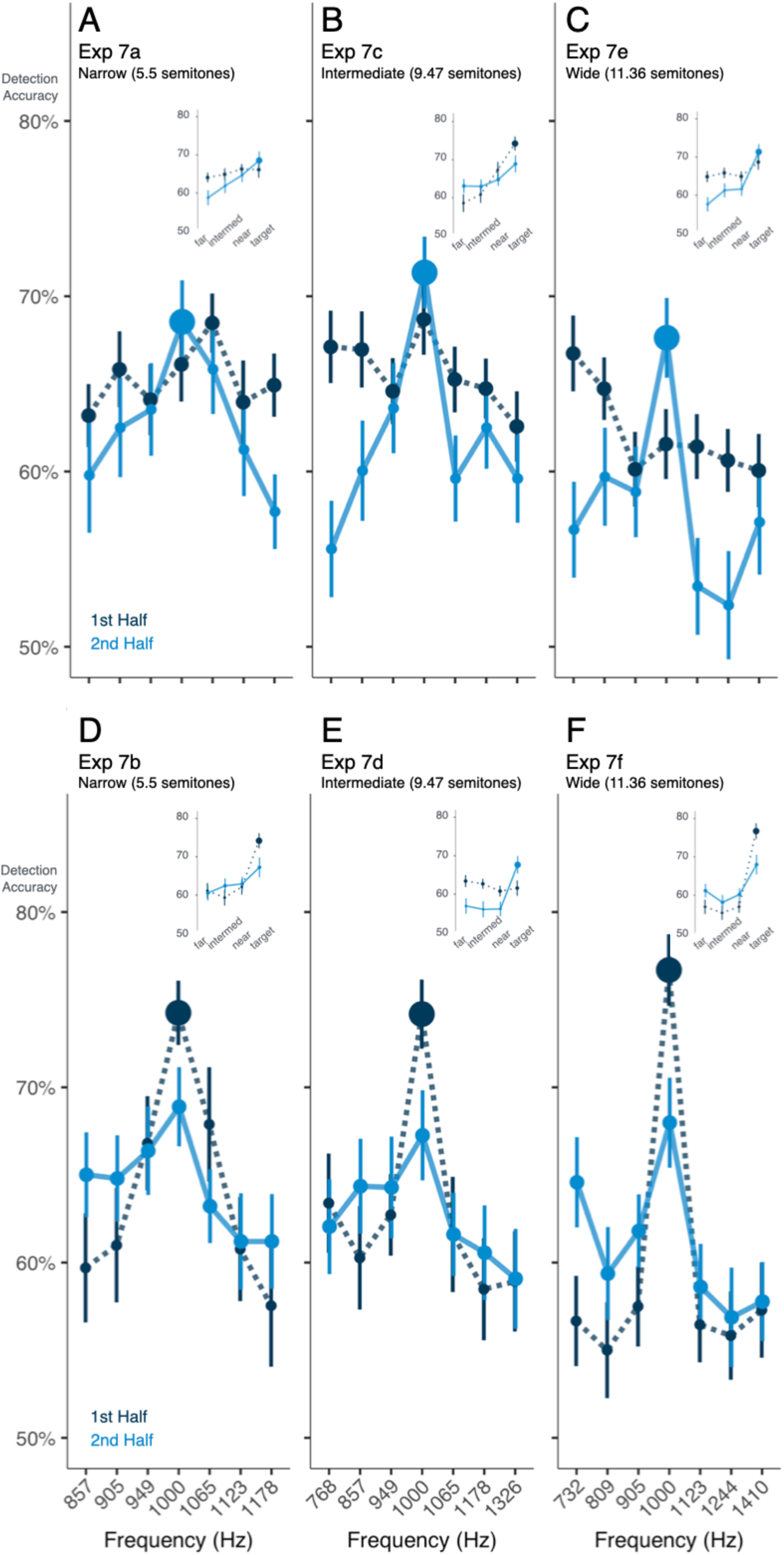
The detailed shape of statistically-driven gain is modulated by range, distribution, and sampling density. See Fig 1c for histograms of distributional regularities. Marker size scales with tone probability. In each panel, the darker color (dotted line) indicates behavior in the first half of the experiment; the lighter color (solid line) indicates behavior in the second half, when distributional regularities shift. Each panel plots mean detection accuracy as a function of acoustic frequency. Error bars indicate standard error of the mean. The top row shows Exp 7a,c,e for which the equiprobable distribution preceded the unimodal distribution. The bottom row shows Exp 7b,d,f for which a unimodal distribution preceded the switch to an equiprobable distribution. Panels **(A)** and **(D)** plot the narrow distribution (5.5 semitone range), Panels **(B)** and **(E)** plot the intermediate distribution (9.47 semitone range), and Panels **(C)** and **(F)** plot the wide distribution (11.36 semitone range). In each panel, the insets show detection accuracy for the high-probability tone (in the unimodal half of the experiment) and equiprobable low-probability tones near, intermediate, and far from the high-probability 1000-Hz tone.

The introduction of the unimodal distribution differentially affects detection, depending on distance of tones from 1000 Hz (ANOVA, p = 1.622 x 10^-11^, η^2^ = 0.030). When 1000 Hz shifts from equiprobable (14.3%) to highly probable (71.4%), there is a small but reliable *increase* in detection accuracy (ANOVA, p = 0.002, η^2^ = 0.026). It is notable that this five-fold increase in probability (and ∼16-fold increase in relative probability compared to low-probability frequencies) only confers an average 3.7% detection improvement. This mild enhancement is not significantly influenced by the range of frequencies (ANOVA, p = 0.365, η^2^ = 0.005). Examining the off-center frequencies that drop in probability (14.3% to 4.8%) upon introduction of a unimodal distribution, we observe a significant *decrease* in detection accuracy of 4.7% (ANOVA, p = 4.798 x 10^-9^, η^2^ = 0.051), the magnitude of which does not differ significantly across range (ANOVA, p = 0.337, η^2^ = 0.003). In brief, when probabilities switch from equiprobable to unimodal we observe a modest increase in detection accuracy for the center frequency that increased in probability and a decrease in detection accuracy for the off-center frequencies that decreased in probability.

Turning next to Exp 7b,d,f (**Fig 6d-f**), what happens when initial experience with a unimodal distribution shifts mid-study to equiprobable presentation? As now expected from prior results, detection of the high-probability mode of a unimodal distribution is considerably more accurate than detection of improbable frequencies (linear contrast, p = 5.198 x 10^-30^, Cohen’s d = 1.332, **Fig 7d-f**). Detection of low-probability frequencies is impacted by proximity to the high-probability center frequency (ANOVA, p = 0.010, η^2^ = 0.016); accuracy is higher for frequencies nearest the high-probability center frequency compared those at middle (p = 0.023, Cohen’s d = 0.269) or far frequencies (p = 0.023, Cohen’s d = 0.274, both contrasts Holm-corrected). However, the relatively preserved detection accuracy for tones near the high-probability frequency compared to those is observed only in Exp 7b for the narrow range (near vs. middle, p = 0.014, Cohen’s d = 0.512, near vs. far, p = 1.161 x 10^-3^, Cohen’s d = 0.697, all linear contrasts Bonferroni-corrected). It is noteworthy that the tones sampling narrow distributions remain highly differentiable at ∼8x larger than typical just-noticeable frequency differences.

**Figure 7.**
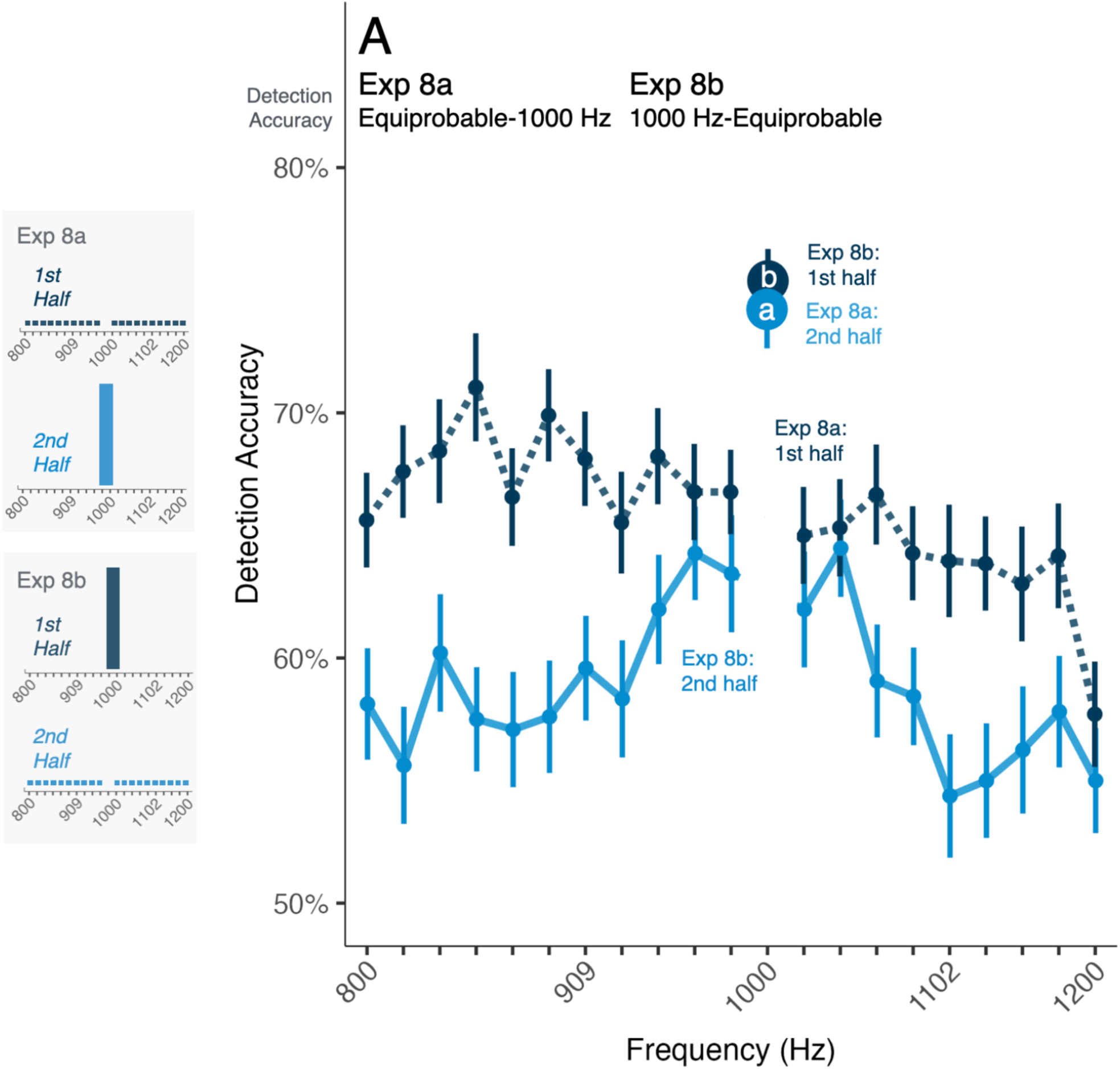
Experience with a single-frequency point distribution results in suppressive reduction in perceptual sensitivity to other frequencies. Exp 8 makes a critical test of whether the gain characterized in the prior experiments involves enhancement of the high-probability frequency, suppression of low-probability frequencies, or a combination of enhancement and suppression. The histograms to the left show distributional regularities for Exp 8a and Exp 8b. Marker size scales with tone probability. Mean detection accuracy is shown as a function of acoustic frequency, with standard error of the mean indicated by error bars. In Exp 8a (dark blue, dashed line), detection trials included 20 equiprobable tones (800-1200 Hz, excluding 1000 Hz) in the first half of the study. In the second half, tones were exclusively 1000 Hz. In Exp 8b (light blue, solid line) the first half of the study involved only 1000 Hz whereas the second half shifted to 20 equiprobable frequencies (800-1200 Hz, excluding 1000 Hz). The inset shows detection in the context of equiprobable distributions for each experiment, as a function of distance from 1000 Hz. Note that detection is somewhat ‘rescued’ around 1000 Hz and that detection of frequencies distant from 1000 Hz is suppressed in Exp 8b relative to Exp 8a.

The effects on detection of proximity to the high-probability 1000 Hz mode are modulated by the switch to an equiprobable distribution (ANOVA, p = 3.279 x 10^-11^, η^2^ = 0.023). We observe a continued, but smaller, detection advantage for the formerly-high-probability center frequency compared to formerly improbable frequencies (linear contrast, p = 1.194 x 10^-8^, Cohen’s d = 0.552). This change is driven by a *decrease* (difference of 7.1%, ANOVA, p = 1.137 x 10^-12^, η^2^ = 0.085) in detection accuracy for the center frequency as it becomes 5 times less probable, as well as a smaller (difference of ∼2%, ANOVA, p = 0.007, η^2^ = 0.006) *increase* in accuracy as off-center frequencies become 3 times more probable; this is potentially compatible with a relative release from suppression. This residual advantage does not vary significantly with distance from the center frequency (ANOVA, p = 0.213, η^2^ = 0.002) or interact with the range of frequencies presented (ANOVA, p = 0.202, η^2^ = 0.004). In sum, there is hysteresis from experience with the unimodal distribution such that the formerly high-probability frequency remains better detected than other frequencies.

Next, we ask if hysteresis is also observed in detection accuracy for 1000 Hz in a unimodal distribution *after* prolonged initial exposure to an equiprobable distribution (second half of Exp 7a,c,e) compared to when the experiment begins with a unimodal distribution (first half of Exp 7b,d,f). We find that pre-exposure to 336 trials of the flat probability distribution diminishes detection rates for the high-probability 1000 Hz tone in the subsequent unimodal distribution by 5.8% relative to when the identical unimodal distribution is encountered first (ANOVA, p = 6.394 x 10^-4^, η^2^ = 0.064). The persistent damping effect of first encountering the equiprobable distribution is not significantly affected by the range of frequencies encountered (ANOVA, p = 0.768, η^2^ = 0.003).

Finally, we aggregate detection data for off-center frequencies across the unimodal conditions from Exp 7a,c,e (when the unimodal distribution was preceded by equiprobable) and Exp 7b,d,f (when it was first) to maximize the power to detect influences of frequency range and distance from the higher-probability center frequency. Frequency range influences detection in unimodal probability distributions (ANOVA, p = 0.005, η^2^ = 0.039). Specifically, a wide frequency range impairs overall off-center detection accuracy, compared to when the frequency range is narrow (post hoc Holm-corrected for following three contrasts, p = 0.006, Cohen’s d = 0.471). The middle frequency range falls in-between and differs significantly from detection in wide (p = 0.037, Cohen’s d = 0.354) but not narrow ranges (p = 0.429, Cohen’s d = 0.118). Moreover, the shape of the drop-off in detection accuracy from the high-probability center frequency is significantly graded only in the narrow frequency range, with a significant difference between the near and mid frequency band conditions (linear contrast, p = 0.015, Cohen’s d = 0.388), and a non-significant decrease between the middle and far frequencies (linear contrast, p = 0.316, Cohen’s d = 0.229).

To summarize Exp 7, we again observe that listeners’ ability to detect a tone in noise is modulated by dynamic changes in statistical distributions. Decreases in probability are met with diminished detection and increases in probability improve detection. However, as we previously observed, the degree of proximity to a more-probable center frequency in unimodal distributions partially rescues detectability of low-probability frequencies. The impact of distributional learning on detection reflects both the probability distribution and the range over which it is defined.

### Experience with a single-frequency point distribution results in suppressive reduction in perceptual sensitivity to other frequencies

The prior experiments leave open the possibility that perceptual interactions across adjacent trials may account for the graded impact on detection, for example through spectrally contrastive influences among tones with different frequencies (Holt, 2005). Exp 8 makes a critical test of whether patterns of relative gain, characterized in the prior experiments, involves enhancement of the high-probability frequency, suppression of low-probability frequencies, or a combination of enhancement and suppression.

To do so, Exp 8 establishes a context in which participants detect *only* 1000 Hz tones in noise, or an equiprobable distribution of 20 tones finely sampling frequency between 800-1200 Hz that *does not include* 1000 Hz (**Fig 1c**). In Exp 8a, the first 320 trials involve 20 different equiprobable (6.25%) tone frequencies (35-cent intervals from 800-1200 Hz, excluding 1000 Hz) and the second 320 trials present exclusively 1000 Hz tones (100% probability). Exp 8b begins with 320 1000-Hz trials, then transitions to the 20-frequency equiprobable distribution (excluding 1000 Hz) across 320 trials. Excluding 1000 Hz from the stimulus set provides a control for possible perceptual interactions across adjacent trials that may have an influence and establishes a baseline against which to evaluate evidence of enhancement and suppression.

We first ask whether the consistent experience with 1000 Hz in the first half of Exp 8b yields accumulating detection accuracy improvements (**Fig 7b**). It does not: accuracy in the first quarter of trials (first half of the first half) is 78% (aligned with expectations from listener-specific thresholding) then decreases slightly to plateau at 75% for the remaining trials in the first half of the study (ANOVA, p = 0.015, η^2^ = 0.068). Similarly, neither Exp 1a (ANOVA, p = 0.210, η^2^ = 0.050) or Exp 8a (ANOVA, p = 0.451, η^2^ = 0.050) exhibit improved detection across a block of trials with only 1000 Hz tones. There is a similar initial detection decrement of ∼5% across the first quarter of the 20-equiprobable-frequency trials of Exp 8a with no further change (ANOVA, p = 9.669 x 10^-6^, η^2^ = 0.049). This same pattern emerges in the initial equiprobable blocks of Exp 7a,c,e (ANOVA, p = 1.375 x 10^-5^, η^2^ = 0.058). Detection accuracy for equiprobable distributions that are experienced in the first half of a study does not differ over experiments (ANOVA, Exp 5a, 7a,c,e, and 8a; p = 0.387, η^2^ = 0.024).

Turning next to the nature of the gain, we first examine whether initial experience with the 20-tone equiprobable distribution in Exp 8a (which does not include 1000 Hz) impacts subsequent detection in the 1000-Hz-only block (**Fig 7a**). It does not: detection of 1000 Hz in the second half of Exp 8a did not differ from either Exp 1a (Tukey-corrected post hoc contrast, p = 0.315, Cohen’s d = 0.333) or the first half of Exp 8b (Tukey-corrected p = 0.837, Cohen’s d = 0.104), each of which involved blocks of trials with only 1000 Hz at the beginning of the study.

In contrast, massed exposure to 1000 Hz in the first half of Exp 8b drives a dramatic, long-lasting, and frequency-specific detection decrement for the subsequently encountered 20 equiprobable frequencies, as compared to detection across equiprobable frequencies in Exp 8a (ANOVA, interaction of Distance-from-1000-Hz x Exp, p = 2.618 x 10^-4^, η^2^ = 0.007). Specifically, as shown in **Fig 7b**, detection of frequencies at far (2 to 3.9 semitones) and intermediate (1 to 2 semitones) distances from 1000 Hz were detected much less accurately after massed experienced with 1000 Hz (Exp 8b, linear contrasts; far: p = 2.427 x 10^-3^, Cohen’s d = 0.330; intermediate: p = 9.784 x 10^-4^, Cohen’s d = 0.346), compared to equiprobable presentation at the beginning of the study (Exp 8a). This suppressive effect was rescued by proximity to the now-absent 1000 Hz in the second half of Exp 8b, with frequencies within about a semitone from 1000 Hz eliciting detection accuracies roughly on par with those from Exp 8a (p = 0.332, Cohen’s d = 0.311). Thus, a half-hour of 1000-Hz exposure induces a lasting attentional filter that impacts the ability to detect frequencies varying from 800-1200 Hz, even though 1000 Hz was never again encountered.

One might expect that any initial learning across the 1000-Hz-only distribution would be overwhelmed by the mid-study shift to the high-uncertainty 20-frequency equiprobable distribution. However, we see the opposite: across the second half of Exp 8b, there is no significant change in overall detection accuracy (ANOVA p = 0.165, η^2^ = 0.008), nor any change across time in relative accuracy of detection across frequencies (ANOVA, p = 0.568, η^2^ = 0.006). The large advantage for detection of frequencies near 1000 Hz compared to intermediate and far frequencies persists to the final 80 trials of Exp 8b (linear contrast, p = 0.002, Cohen’s d = 0.257). This effect is further evidenced by comparing the second half of Exp 8b with the first half of Exp 8a. Here, there is strong suppression of frequencies at far and intermediate distances from 1000 Hz in Exp 8b compared to detection of the same frequencies in the equiprobable half of Exp 8a. As for the within-experiment comparison, this difference is observed through the entirety of the second half of the study, again extending even to the last quarter of trials (linear contrast, p = 0.008, Cohen’s d = 0.440). The absence of 1000 Hz over this period rules out the possibility that trial-wise perceptual interactions or the experience of a relative probability difference for a particular frequency were strong contributors to the hysteresis observed in Exp 5 and Exp 7. See Fig S2.

## Discussion

Is perception guided toward what we expect, or by what surprises us? Here, across 29 experiments, we examine two perceptual tasks for which distributional regularities accumulate over a task-irrelevant dimension without instruction, directed attention, or feedback. We find that distributional learning drives dynamic shifts in perception across tasks. It affects sound detection: a faint tone of a particular frequency is better detected in noise if it occurs frequently than if it occurs rarely. Distributional learning also affects duration decisions, which are faster for tones that possess a frequency that has been more common in recent experience. Across tasks, our results converge to indicate that effects of expectation are largely driven by suppression of less-probable, unexpected stimuli that are, to a lesser degree, supported by modest enhancement of highly probable stimuli.

Our examination of expectation built across distributions (rather than dichotomous probabilities) affords a wider vantage point for understanding how perceptual gain is modulated by expectation. Our results reveal an influence on perception that is graded as a function of the distribution mode, the range of the distribution, and the position of a stimulus within the distribution. The detailed shape of the distribution is important, as well, as shown by the bimodal profile of tone detection evoked by a bimodal frequency distribution. Strikingly, equally probable rare events are perceived differently as a function of their perceptual distance from the distribution mode(s). Decades ago, Greenberg and Larkin (1968) examined tone detection in a similar paradigm (albeit with overt instructions about tone probability instead of distributional learning) and interpreted the graded gain to be indicative of a frequency-selective attentional filter situated at the high-probability mode with increasingly suppressive sidebands with greater distance from the mode. Indeed, in the time since there has been sustained interest (e.g., Summerfield & Egner, 2009; Zivony & Eimer, 2024) in isolating the influence of *expectation* - operationalized by manipulating the probability of stimuli – from *attention* – defined according to the utility or relevance of these stimuli to a task (Summerfield & de Lange, 2014; Kok et al., 2012). Under these definitions, the present tasks are attention-neutral and involve manipulations of *expectation* only. Yet, our results suggest that expectation built across distributional learning establishes a selection filter that impacts how (and whether) subsequent stimuli are perceived. Whether this is described as a dimension-selective attentional filter (as proposed by Greenberg & Larkin, 1968) or more neutrally as an experience-driven predictive filter, the present results are distinct from manipulations of task utility or relevance that have been previously attributed to attention interacting with expectation (Zivony & Eimer, 2024; Rungratsameetaweemana & Serences, 2019).

In the time domain, the influence of distributional learning on perception is persistent: effects of a unimodal distribution provoke lasting influence with a continued advantage for tones that were previously probable and a lasting disadvantage for the tones that were previously improbable, even after exposure to a uniform distribution. Even so, there remains sensitivity to volatile distribution changes with both detection and perceptual decisions dynamically adjusting when dichotomous probabilities flip. Future work will be needed to resolve the interpretive tension between the rapid adjustment we observe across changing dichotomous probabilities in Exp 3 and Exp 4 versus the lingering influence of bimodal (Exp 5,6,7) and point (Exp 8) distributions. Candidate contributors include the magnitude of differences in stimulus probabilities, dichotomous versus more fully sampled distributions, lower information conveyance by uniform distributions, and the relative volatility experienced across a listening session. The present paradigms provide a basis for further discovery, with implications for ‘stubborn predictions’ examined in other literatures (Yon et al., 2023).

The impact of these distributional regularities on perception is evident for both detection and perceptual decisions, emphasizing the breadth of influence of distributional learning across distinct task demands (Fritsche, Mostert, & de Lange, 2017). Even so, detection provides a unique window through which to observe effects of distributional learning and resulting expectations, as it has a natural baseline set by individuals’ thresholds. The detection results make it especially clear that the net impact of distributional learning is to prioritize the high-probability distribution mode - not by enhancing detectability of the expected stimulus, but instead by suppressing detectability of rare, unexpected stimuli. We observe this repeatedly across experiments. Despite considerable headroom for detection accuracy to improve in the context of a threshold set at ∼79% accuracy we do not observe substantial enhancement of detection of the high-probability tone. Indeed, in the original Greenberg and Larkin (1968) study, exposure to tens of thousands of trials of a high-probability frequency did not enhance detection above the initially established perceptual threshold. This lack of enhancement at the mode is somewhat surprising given the literature on perceptual learning (Amitay, Zhang, Jones, & Moore, 2014; Watanabe & Sasaki, 2015; Wright, Wilson, & Sabin, 2010), where intensive practice with attentionally-demanding perceptual paradigms can drive improved detection. But, in contrast to most perceptual learning approaches, the influences we observe accrue across a task-irrelevant perceptual dimension, without directed attention, reward, or feedback.

It would seem inefficient for a system to track distributional regularities irrelevant to the task at hand. However, ‘optimal’ selectivity to a task-relevant dimension may not always be adaptive for perception: in natural environments with shifting demands, it may be effective to ‘keep an ear out’ by tracking evolving regularities with potential utility for future behavior (Heilbron & Chait, 2017). Moreover, the reduction in sensitivity to subsequently encountered frequencies that we observe following massed exposure to a single frequency would seem to be a maladaptive loss of perceptual sensitivity. Instead, it may reflect gain mechanisms that suppress sensitivity to region(s) along a perceptual dimension that are less likely to be encountered. In the sense that one cannot be surprised by something if one is not sure it has occurred (Press et al., 2020), the suppressive effects we observe for low-probability stimuli distant from a distribution mode are substantial enough that these stimuli would seem to be less likely to contribute to subsequent distributional learning. Distributional effects on perception thus may have the potential to snowball, exaggerating regularities relative to the true distribution of events.

Our results emphasize that layered histories of experience with distributional regularities impact behavior. For example, unimodal distributions have lingering effects even after a switch to equiprobable stimulus presentation. At a longer timescale, we observe a consistent frequency-duration bias in our perceptual decision experiments. The effect is persistent across decision experiments (even when only two frequencies were present) and appears to be associated with the ordinal position of frequencies in the distribution range rather than absolute frequency. Although acoustic frequency and duration would seem to be good candidates for orthogonal acoustic input dimensions – and indeed, older studies had suggested this (Allan & Kristofferson, 1974; Woods, Sorkin, & Boggs, 1979) – the ubiquity of interactions between acoustic dimensions is seen clearly in auditory category learning studies in which rotating the sampling of acoustic category exemplars in an ostensibly orthogonal acoustic space produces radically different learning outcomes due to prior expectations about the relationship between the dimensions (Roark & Holt, 2022; Bröker et al., 2024; Bröker, Plaut & Holt, 2024).

Life-long exposure to the distributional statistics of natural sound environments may drive at least some of the ubiquitous bias to perceive relatively lower frequencies as longer, and relatively higher frequencies as shorter (Fiser, Berkes, Orbán, & Lengyel, 2010; Berkes, Orbán, Lengyel, & Fiser, 2011). Pinning down the etiology of this endogenous bias will be challenging, as multiple environmental and acoustic factors may contribute. From different decay characteristics for struck strings on the piano (undamped bass notes decay much more slowly than treble notes; Fletcher, Blackham & Stratton, 1962) to the longer reverberance for lower versus higher frequencies (Backus, 1977) there are complex, and likely consistent, regularities across acoustic frequency and duration that individuals may learn about over a lifetime of listening. Just as important, there may be sensory contributions such that the longer duration of cochlear filters at lower frequencies may facilitate identification of lower-frequency tones as long duration, consistent with the bias we observe across experiments (Patterson, 1987). For example, an auditory filter centered at 800 Hz is expected to have a longer duration than a filter centered at 1200 Hz. Thus, longer-lasting peripheral excitation at lower frequencies may facilitate identification of lower-frequency tones as long duration.

Sensory factors may play a role in the subtle influence across frequencies observed for tone detection, as well. As noted above, the cochlea’s limited ability to resolve stimuli creates critical bands within which it is difficult to distinguish frequencies (Fletcher, 1940). As described above, the ‘rescue’ of detection accuracy for low probability tone frequencies that lie closer to the distribution mode may partly reflect limitations of frequency resolution within a critical band. Similarly, changes in the width of the critical band across our 800-1200 Hz frequency range may account for the subtle, consistent tendency for detection to be less accurate at higher frequencies (Fig 2, 5, 6). Yet, beyond sensory influences, our manipulation of distributions (instead of binary probabilities) helps to reveal that other factors are at play. For example, both the distribution range and the position of the high probability tone with the distribution range influence detection.

The present results are potentially informed by rich literatures studying neural response across stimuli that vary in probability. Repeated exposure to a stimulus changes neural firing patterns in visual (Schoups, Vogels, Qian, & Orban, 2001) and auditory (Khouri & Nelken, 2015) cortex. Two neural phenomena - the mismatch negativity (MMN, Naatanen et al., 1978), and stimulus specific adaptation (SSA, Ulanovsky et al., 2004) – are extensively studied in the auditory domain using an ’oddball’ paradigm in which common and rare stimuli are intermixed in a sequence. This probability manipulation reveals exaggerated neural response to low-probability sounds, seeming to run counter to the principally suppressive behavioral effects we observe for low-probability tones. However, these neural phenomena can be observed in active freely moving states (Polterovich et al., 2018) as well as under anesthesia (Yaron et al., 2012) and in disordered consciousness (Bekinschtein et al., 2009); we do not yet have a strong understanding of how they relate to auditory behavior. There is much more to understand in relating exaggerated neural response to low-probability sounds with slower decisions and less accurate detection. Nonetheless, these neurobiological literatures support the possibility that effects of distributional learning can emerge robustly in primary auditory cortex, and possibly even auditory midbrain (Malmierca et al., 2009; Duque et al., 2012; Cruces-Solís et al., 2018).

Schröger and Wolf (1998), who pioneered the duration-decision task we use here, argued from human electroencephalography results that, at least for perceptual decisions, effects may arise from a memory-based mechanism that detects deviance from expectations and orients attention to the rare stimulus frequency. This leaves fewer resources for perceptual decision making, slowing response times. However, in a case of convergent experimental design, Mondor and Bregman (1994) used a very similar duration-decision paradigm to argue that the reaction time advantage for probable or cued frequencies instead arises from allocation of selective attention to the probable (not the improbable) frequency. This echoes the interpretational challenges of the larger literature on expectation and attention effects (see Introduction) and is mirrored, as well, in literatures attempting to resolve patterns of behavioral repetition priming and neural repetition suppression (McMahon & Olson, 2007; Feuerriegel, Vogels, & Kovács, 2021).

Organisms as diverse as humans and honeybees are exquisitely sensitive to patterns that unfold across sensory input. We find that distributional learning affects fundamental aspects of perception: the very ability to detect whether a stimulus is present and to make a simple judgment about it. Listeners rapidly and implicitly apprehend distributional regularities of how often stimuli occur, even when the regularities emerge across sensory dimensions irrelevant to the task at hand. This statistical learning across input distributions arises rapidly even in the context of statistically dynamic contexts and has a substantial and lasting influence on perception driven by collaborative influences of sensory processing, distributional learning, and selective attention in sculpting a gain function involving modest enhancement and robust suppression.

## Materials and Methods

Experiment materials, code, and analyses can be found at https://osf.io/xdgnw/.

### Participants

Participants (ages 18-35 yrs) were recruited online and compensated via Prolific.co (Damer & Bradley, 2014). All self-reported normal hearing. **Table S1** provides experiment-wise demographic details. Based on power analyses of pilot data collected using the same tasks, we targeted recruitment of 30 participants/experiment.

### Stimuli

Sinewave tones and white noise were generated in the lossless FLAC format using the Sound eXchange sound processing software (SoX, http://sox.sourceforge.net/) at 44.1kHz and 16-bit precision.

### Procedure

All experiments were conducted online following best-practices described by Zhao et al. (2022) using PsychoPy (2022.1.2, pavlovia.org) for tone-in-noise detection experiments and Gorilla (Anwyl-Irvine et al., 2020) for duration-decision experiments. Online participants used the Chrome browser on their own laptop or desktop computer (no smartphones or tablets) with a brief listening test assuring headphone compliance (Milne et al., 2020). **Fig 1** illustrates the trial structure for each task. Table S2 provides experiment-level details.

#### Tone-in-Noise Detection Task

Continuous white noise commenced +40 dB relative to the level just detectable over participants’ own computer and headphones, as determined by a brief system-calibration procedure (Zhao et al., 2022). Adaptive thresholding commenced with the onset of a 300-sec white noise (200-ms cosine amplitude onset/offset ramps) that looped continuously through the end of the study. Adaptive thresholding entailed detecting a 250-ms (10-ms cosine onset/offset ramps), 1000-Hz sinewave tone (1080-Hz in Exp 1f) in a three-interval forced choice task (**Fig 1a**). The first 6 trials served as practice, with feedback and -13.75 dB SNR. Thereafter, there was no feedback across three 40-trial adaptive thresholding runs. Each run began at -13.75 dB SNR with tone intensity decreasing 1.5 dB after each correct detection until the SNR reached -19.75 dB, or until an incorrect response. Subsequently, tone intensity decreased -.75 dB after three correct responses and increased +.75 dB after each incorrect response. Threshold tone-in-noise detection was computed as the mean of the mode tone intensity across the three runs (Zhao et al. 2022) which estimates threshold at 79.4% correct detection (Levitt, 1971).

Adaptive thresholding established a by-participant threshold tone intensity for the tone-in-noise experiment. The first experiment block was practice, with -13.75 dB SNR, feedback, and tone frequencies that matched the initial experiment distributional regularity (**Fig 1a**). After practice, tone intensity was set to -.75 dB relative to the threshold estimate for the remainder of the experiment. Participants reported which of two intervals contained the tone (**Fig 1a**). Participants were not informed about the task-irrelevant distributional regularities across acoustic frequency (**Fig 1c**). The entire protocol took about 30 minutes, except in experiments with double the trials (see **Table S2**). We report mean detection accuracy.

#### Duration-Decision Task

Each trial involved a single sinewave tone presented in quiet at a comfortable level. Tones were 50 or 90 ms, with equal probability and random presentation. Participants reported whether the tone was “long” or “short” with a key press and were not instructed about the task-irrelevant distributional regularities across acoustic frequency (**Fig 1b**). Each experiment began with a practice block involving feedback and a distributional regularity that mirrored the main experiment. There was no feedback for the remainder of the experiment. **Table S2** provides experiment-wise details. The entire protocol took about 30 minutes, except in experiments with double the trials. Analyses focused on decision response time, measured from tone offset to response. Trials for which response time was shorter than 300 ms or longer than 1500 ms (non-inclusive) were excluded from analyses (see **Table S1** for percent of trials excluded).

### Approach to Analysis

Data were preprocessed using JMP Pro 17.0.0, and statistical analyses were conducted in JASP (JASP team, Amsterdam, Netherlands, 10/19/22, version 0.16.4). We report Greenhouse-Geisser corrected degrees of freedom and *p* values for ANOVAs for which the assumption of sphericity was violated, as determined by a Mauchly test. Multiple comparison correction for linear contrasts was carried out using Bonferroni correction, and for post-hoc tests using Holm correction. Study-wise analysis details are provided in **Table S3**.

## Acknowledgements and Funding Sources

The work was supported by grants from National Institutes of Health (R01DC004674) and the National Science Foundation (SBE/BCS 2414066, 2420979) to LLH and FD, an institutional training grant from the National Institute of General Medical Sciences to LLH that supported AL (T32GM081760), and an individual NRSA from the National Institute on Deafness and Communication Disorders to SL (F32DC020625) under the mentorship of LLH and FD. The work reported here comprised a chapter of AL’s 2024 PhD dissertation (Carnegie Mellon University). Christi Gomez, Erin Smith, and Gracee Swatsworth contributed to the study. Drs. Clare Press, Daniel Yon, and Alain de Chevigné provided very helpful comments on a manuscript draft.

