## Supplemental Information for "Statistical learning dynamically shapes auditory perception"

### Supporting Information for
### Distributional learning drives statistical deafening

Sahil Luthra[1], Austin Luor[1], Adam T. Tierney, Frederic Dick, Lori L. Holt

Professor Lori L. Holt


[1] equal contributions

**This PDF file includes:**

Tables S1 to S3
Figures S1 to S2

#### Table S1. Participant Demographics

*Note that age and gender reflect only those participants included in analyses. For the tone-in-noise detection task, exclusion was by participant and not by trial. Exclusion criteria for the tone-in-noise detection task included: (1) very high tone-in-noise thresholds (-14.5 dB SNR or higher) consistent with extremely poor performance and the possibility of hearing loss, inattention or other challenges to participation; (2) individuals who reported not hearing the continuous white noise masker for the entirety of the experiment; (3) participants with very low thresholds (-26.5 dB SNR or lower) that led us to suspect errors in masking noise presentation. Exclusion criteria for the duration decision task included: (1) performance on no-feedback trials that was at or below chance, on average. At-chance performance was determined based on permutation testing with 10,000 permutations and an alpha level of 0.05.* ***Table 2*** *indicates at-chance performance levels for each duration decision experiment. In addition, incorrect decisions and decisions that were made fewer than 300 ms after stimulus onset or 1500 ms after stimulus onset were excluded on a by-trial basis.*

| Experiment | Age (SD) | # Participants (Gender) | # Excluded Participants | % Trial Loss (Duration Decision Only) |
|---|---|---|---|---|
| **Exp 1a** | 24.5 (3.8) | N = 28 (5 female, 22 male, 1 non-binary) | 2 | — |
| **Exp 1b** | 24.7 (4.0) | N = 28 (12 female, 16 male) | 2 | — |
| **Exp 1c** | 26.7 (5.2) | N = 29 (16 female, 13 male) | 1 | — |
| **Exp 1d** | 25.5 (3.7) | N = 27 (13 female, 14 male) | 3 | — |
| **Exp 1e** | 24.4 (4.1) | N = 28 (12 female, 16 male) | 4 | — |
| **Exp 1f** | 24.8 (3.9) | N = 30 (13 female, 16 male, 1 non-binary) | 0 | — |
| **Exp 2a** | 23.7 (4.2) | N = 30 (10 female, 20 male) | 0 | 28.11% |
| **Exp 2b** | 23.4 (3.7) | N = 30 (13 female, 17 male) | 0 | 25.48% |
| **Exp 2c** | 25.4 (4.6) | N = 29 (10 female, 19 male) | 1 | 25.16% |
| **Exp 2d** | 25.2 (4.4) | N = 27 (11 female, 16 male) | 3 | 23.84% |
| **Exp 2e** | 25.4 (5.5) | N = 30 (7 female, 23 male) | 0 | 21.97% |
| **Exp 3a** | 24.0 (4.0) | N = 28 (14 female, 14 male) | 3 | — |

| | | | | |
|---|---|---|---|---|
| **Exp 3b** | 23.4<br>(3.6) | N = 30<br>(16 female, 14 male) | 1 | — |
| **Exp 3c** | 24.4<br>(3.7) | N = 28<br>(12 female, 16 male) | 2 | — |
| **Exp 4a** | 23.4<br>(3.6) | N = 31<br>(11 female, 19 male, 1 non-binary) | 0 | 15.32% |
| **Exp 4b** | 24.7<br>(4.6) | N = 29<br>(8 female, 21 male) | 0 | 14.61% |
| **Exp 4c** | 24.6<br>(4.8) | N = 27<br>(10 female, 17 male) | 3 | 15.39% |
| **Exp 5a** | 26.0<br>(5.7) | N = 27<br>(13 female, 14 male) | 3 | — |
| **Exp 5b** | 24.9<br>(3.7) | N = 30<br>(12 female, 18 male) | 0 | — |
| **Exp 6a** | 24.3<br>(4.1) | N = 28<br>(9 female, 17 male, 1 non-binary) | 2 | 25.31% |
| **Exp 6b** | 24.5<br>(4.1) | N = 29<br>(10 female, 18 male, 1 non-binary) | 1 | 22.30% |
| **Exp 7a** | 26.7<br>(4.5) | N = 30<br>(11 female, 17 male, 2 non-binary) | 0 | — |
| **Exp 7b** | 26.0<br>(4.4) | N = 30<br>(7 female, 23 male) | 1 | — |
| **Exp 7c** | 25.4<br>(4.3) | N = 30<br>(5 female, 25 male) | 2 | — |
| **Exp 7d** | 24.3<br>(3.9) | N = 30<br>(10 female, 19 male, 1 non-binary) | 2 | — |
| **Exp 7e** | 24.9<br>(4.8) | N = 29<br>(7 female, 22 male) | 2 | — |
| **Exp 7f** | 24.0<br>(4.1) | N = 30<br>(9 female, 21 male) | 0 | — |
| **Exp 8a** | 24.7<br>(4.3) | N = 60<br>(19 female, 39 male, 1 non-binary,<br>1 prefer not to answer) | 3 | — |
| **Exp 8b** | 25.8<br>(4.6) | N = 60<br>(18 female, 41 male, 1 non-binary) | 5 | — |

**Table S2. Summary of Experiment Protocols**

*Detailed protocol for each experiment, according to task, tone frequencies, trials, trials/block, and the number of statistical distribution changes. Chance performance for the duration decision task was determined based on permutation testing with 10,000 permutations and an alpha level of 0.05.*

| Exp | Most probable frequency | # Trials | Trials / Block | When do statistics change? | Chance-level performance (duration decision) |
|---|---|---|---|---|---|
| **Exp 1a** (Tone-in-noise detection) | 1000 Hz (100% of trials) | 320 | 32 | n/a | n/a |
| **Exp 1b** (Tone-in-noise detection) | 920 Hz (75% of trials) | 320 | 32 | n/a | n/a |
| **Exp 1c** (Tone-in-noise detection) | 1000 Hz (75% of trials) | 320 | 32 | n/a | n/a |
| **Exp 1d** (Tone-in-noise detection) | 1080 Hz (75% of trials) | 320 | 32 | n/a | n/a |
| **Exp 1e** (Tone-in-noise detection) | 800 Hz and 1200 Hz (40.625% of trials each) | 320 | 32 | n/a | n/a |
| **Exp 1f** (Tone-in-noise detection) | 800 Hz and 1200 Hz (40.625% of trials each) | 320 | 32 | n/a | n/a |
| **Exp 2a** (Duration decision) | 920 Hz (80% of trials) | 400 | 40 | n/a | 54.0% |
| **Exp 2b** (Duration decision) | 1000 Hz (80% of trials) | 400 | 40 | n/a | 54.0% |
| **Exp c** (Duration decision) | 1080 Hz (80% of trials) | 400 | 40 | n/a | 54.0% |

| | | | | | |
|---|---|---|---|---|---|
| **Exp 2d** (Duration decision) | 800 Hz and 1200 Hz (40.625% of trials each) | 320 | 32 | n/a | 54.7% |
| **Exp 2e** (Duration decision) | 1st half: all frequencies equiprobable 2nd half: 800 Hz and 1200 Hz (42.5% of trials each) | 640 | 40 | After 8 blocks | 53.3% |
| **Exp 3a** (Tone-in-noise detection) | 1000 Hz (75% of trials) | 320 | 32 | n/a | n/a |
| **Exp 3b** (Tone-in-noise detection) | 1155 Hz (75% of trials) | 320 | 32 | n/a | n/a |
| **Exp 3c** (Tone-in-noise detection) | 1000 or 1155 Hz, switching throughout experiment (75% of trials) | 640 | 32 | After every 5 blocks | n/a |
| **Exp 4a** (Duration decision) | 1000 Hz (80% of trials) | 320 | 32 | n/a | 54.7% |
| **Exp 4b** (Duration decision) | 1155 Hz (80% of trials) | 320 | 32 | n/a | 54.7% |
| **Exp 4c** (Duration decision) | 1000 or 1155 Hz, switching throughout experiment (75% of trials) | 640 | 32 | After every 5 blocks | 53.3% |
| **Exp 5a** (Tone-in-noise detection) | 1st half: all frequencies equiprobable 2nd half: 1000 Hz (80%) | 640 | 40 | After 8 blocks | n/a |
| **Exp 5b** (Tone-in-noise detection) | 1st half: 1000 Hz (80%) 2nd half: all frequencies equiprobable | 640 | 40 | After 8 blocks | n/a |
| **Exp 6a** (Duration decision) | 1st half: all frequencies equiprobable 2nd half: 1000 Hz (80%) | 640 | 40 | After 8 blocks | 53.3% |

| | | | | | |
|---|---|---|---|---|---|
| **Exp 6b** (Duration decision) | 1st half: 1000 Hz (80%) 2nd half: all frequencies equiprobable | 640 | 40 | After 8 blocks | 53.3% |
| **Exp 7a** (Tone-in-noise detection) | 1st half: all frequencies (857, 905, 949, 1000, 1065, 1123, 1178 Hz) equiprobable 2nd half: 1000 Hz (71.4%) | 672 | 42 | After 8 blocks | n/a |
| **Exp 7b** (Tone-in-noise detection) | 1st half: 1000 Hz (71.4%) 2nd half: all frequencies (857, 905, 949, 1000, 1065, 1123, 1178 Hz) equiprobable | 672 | 42 | After 8 blocks | n/a |
| **Exp 7c** (Tone-in-noise detection) | 1st half: all frequencies (768, 857, 949, 1000, 1065, 1178, 1326 Hz) equiprobable 2nd half: 1000 Hz (71.4%) | 672 | 42 | After 8 blocks | n/a |
| **Exp 7d** (Tone-in-noise detection) | 1st half: 1000 Hz (71.4%) 2nd half: all frequencies (768, 857, 949, 1000, 1065, 1178, 1326 Hz) equiprobable | 672 | 42 | After 8 blocks | n/a |
| **Exp 7e** (Tone-in-noise detection) | 1st half: all frequencies (732, 809, 905, 1000, 1123, 1244, 1410 Hz) equiprobable 2nd half: 1000 Hz (71.4%) | 672 | 42 | After 8 blocks | n/a |
| **Exp 7f** (Tone-in-noise detection) | 1st half: 1000 Hz (71.4%) 2nd half: all frequencies (732, 809, 905, 1000, 1123, 1244, 1410 Hz) equiprobable | 672 | 42 | After 8 blocks | n/a |
| **Exp 8a** (Tone-in-noise detection) | 1st half: 20 frequencies (range: 800-1200 Hz, 37-cent steps), all equiprobable (5% each) 2nd half: 1000 Hz (100% of trials) | 640 | 40 | After 8 blocks | n/a |

| | | | | | |
|---|---|---|---|---|---|
| **Exp 8b** (Tone-in-noise detection) | 1st half: 1000 Hz (100% of trials)<br>2nd half: 20 frequencies (range: 800-1200 Hz, 37-cent steps), all equiprobable (5% each) | 640 | 40 | After 8 blocks | n/a |

**Table S3. Detailed Analysis Information.**

| Exp | Text from Results | Test | Test in JASP (exp cond filter) | JASP datafile | Notes | Excluded Ss |
|---|---|---|---|---|---|---|
| **1b-f** | Frequency probability strongly modulates tone detection in noise across Exp 1b-f | Frequency x Exp interaction | 1b-f, exclude a (although doesn't matter) | Exp1a-f-TPS-AccSummary240508.jasp | | |
| **1a, 1b-d** | Detection of only 1000 Hz (Exp 1a: 100% probability; average accuracy 77.9%) does not differ from detection of the highest-probability frequency in unimodal distributions (Exp 1b-d: 75% probability; average accuracy 75.3%; p = 0.242). | ANOVA | exclude ProbabilityOfFrequency '40' (TPS dual conditions, e-f) | Exp1a-f-MostProbableFreqAcc.jasp | | |
| **1b-d, 1e-f, 1a** | But detection of the 40%-probable frequencies in bimodal distributions is lower than when a single frequency is 80% or 100% probable (Exp 1e-f: 40.6% probability; average accuracy 70.3%; p = 0.006 versus Exp 1b-d, p = 0.003 versus Exp 1a). | ANOVA | exclude ProbabilityOfFrequency '100' (TPS 1000) OR '80' (all others) | Exp1a-f-MostProbableFreqAcc.jasp | | |
| **1b-d** | Proximity to the high probability tone also influences detection (**Fig 3a**). The low-probability frequencies of Exp 1b-d share the same probability, yet those closer to a high-probability frequency are better detected than those further away (p = 0.014). | RM ANOVA, Main effect of DistFromProbFreq | none | Exp1b-c-d-TPS-byDistFromHighProbFreq-240508.jasp | | |
| **1c** | When the high-probability frequency is centered in the range of frequencies defining the distribution, this graded detection accuracy difference is symmetric (far < near to high-probability frequency, p = 0.004). | RM ANOVA, Main effect of DistFromProbFreq | DistanceFromHighProbabilityFrequency,FilterByExpt, Filter out b and d | Exp1b-c-d-TPS-byDistFromHighProbFreq-240508.jasp | | |

| | | | | | | |
|---|---|---|---|---|---|---|
| **1b, 1d** | When the high-probability frequency is nearer to the distribution edge (Exp 1b and Exp 1d), there is an asymmetric detection curve (p = 0.015): a sharp detection decrement toward the distribution edge is contrasted with a more gradual 'ski slope' decrement toward the middle of the frequency range. | Main effect, DistAccordingToPosInRange | none | Exp1b-d-TPS-EdgeMiddleNearMidFar.jasp | Edge frequency accuracy is < Middle (p = 0.016), and Far < Middle (p = 0.038). | |
| **1e** | Exp 1e shows that a bimodal probability distribution with higher-probability (40.6%) frequencies at the edge of the spectral range (800 and 1200 Hz) induces a 'dual spotlight' across the frequency dimension. Listeners detect the higher-probability tones more accurately than neighboring low-probability tones (920 and 1080 Hz, $p = 3.451 \times 10^{-7}$) as well as the middle 1000 Hz tone (p = 0.036). | SingleConditionANOVA Exp1e OR f With contrasts, Freq effect; linear contrasts Bonferroni corrected (e.g., p-values x # of contrasts = 3) | filter out 'f' | Exp1e-f-TPS-AccSummary240508.jasp | | |
| **1f** | Exp 1f falsifies this hypothesis. Changing the initial threshold-setting frequency to 1080 Hz elicits a similar "W" profile and, importantly, replicates the overall 'dual spotlight' at 800 and 1200 Hz ($p = 8.52 \times 10^{-11}$, **Fig 3b**). | SingleConditionANOVA Exp1e OR f With contrasts, Freq effect; linear contrasts Bonferroni corrected (e.g., p-values x # of contrasts = 3) | filter out 'e' | Exp1e-f-TPS-AccSummary240508.jasp | Reported p-value is for the Bonferroni-corrected contrast of 800 & 1200 vs 920 & 1080 (contrast 1) | |
| **2a-c** | Across Exp 2a-c, the probability of a tone's *frequency* significantly impacts the speed of duration decisions ($p = 7.62 \times 10^{-7}$, **Fig 4**). | Main effect of frequency, RM ANOVA | none | Exp2a-c-PSID-FiltRTsByFreq.jasp | | (Excluded 1 subject: 5304786 from 1080Bias) |
| **2a-c** | Response times (RTs) vary are faster for tones with high, compared to low, probability frequencies ($p = 1.445 \times 10^{-21}$). | Linear contrast of HighPro vs Far+Near | none | Exp2abc-AllConds-FINAL-RT-Filt300-1500ms-FreqXDur.jasp | | (Excluded 1 subject: 5304786 from 1080Bias) |

| | | | | | | |
|---|---|---|---|---|---|---|
| **2a-c** | RTs to the most probable frequency are faster than the adjacent low-probability frequencies ($p = 5.222 \times 10^{-11}$), which are in turn faster than frequencies furthest away from the high-probability frequency ($p = 4.19 \times 10^{-6}$). | Post-hoc comparisons table, holm-corrected | none | Exp2abc-AllConds-FINAL-RT-Filt300-1500ms-FreqXDur.jasp | Holm's corrected posthoc table | (Excluded 1 subject: 5304786 from 1080Bias) |
| **2a-c** | (These patterns hold true for each Exp 2a-c study, $p < .05$ Holm-corrected). | | filter by each condition separately | Exp2abc-AllConds-FINAL-RT-Filt300-1500ms-FreqXDur.jasp | Holm's corrected posthoc table | (Excluded 1 subject: 5304786 from 1080Bias) |
| **2d** | However, unlike the dual spotlight for tone detection in Exp 1e-f, there is no significant RT advantage for making duration decisions about the higher-probability 800 and 1200 Hz tones in Exp 2d (**Fig 4**; $p = 0.615$). | RM-ANOVA, main effect of frequency | none | Exp2d-PSID-Bimodal-RT-FreqXDuration.jasp | | (Excluded 3 subjects: 6492029, 6492881, 6491877) |
| **2e** | Indeed, decision RTs are *longer* for 800 Hz and 1200 Hz compared to other frequencies ($p = 0.031$) when tone frequencies are equiprobable in the first half of trials. | | no filters | Exp2e-PSID-EquiToBimodal-RT-FreqXDuration.jasp | note in general that Greenhouse-Geisser always used | |
| **2e** | Investigating this, we observe a *novel frequency-duration perceptual bias*: duration decisions for lower-frequency tones (800, 920 Hz) are more accurate and faster for long (90 ms) compared to short (50 ms) tones whereas those for the highest frequency tone (1200 Hz) are more accurate and faster for short compared to long tones (**Fig S1**; Frequency x Duration interaction, RT: $p = 0.003$, | RM-ANOVA, interaction of frequency and duration "RM-ANOVA on RTs with duration as factor" | | Exp2e-PSID-EquiToBimodal-RT-FreqXDuration.jasp | G-G | |
| **2e** | ….Accuracy (Acc): p = 3.738 x 10-5) | Repeated Measures ANOVA, Frequency x Duration interaction | none | Exp2e-PSID-EquiToBimodal-Acc-FreqXDuration.jasp | G-G | |

| | | | | | | |
|---|---|---|---|---|---|---|
| **2d** | This longer-lower/shorter-higher bias is mirrored qualitatively in Exp 2d (**Fig S1**; $p > 0.05$) | Repeated Measures ANOVA, Frequency x Duration interaction | none | Exp2d-PSID-Bimodal-RT-FreqXDuration.jasp | | (Excluded 3 subjects: 6492029, 6492881, 6491877) |
| **2a-c** | When we inspect the data from Exp 2a-c (**Fig S1**), we also observe the longer-lower/shorter-higher bias in the context of the unimodal distributions (Frequency x Duration interaction, RT: $p = 3.968 \times 10^{-6}$; | Repeated Measures ANOVA, Frequency x Duration interaction | none | Expt2abc-AllConds-FINAL-RT-Filt300-1500ms-FreqXDur.jasp | | (Excluded 1 subject: 5304786 from 1080Bias) |
| **2a-c** | Acc: $p = 0.003$). | Repeated Measures ANOVA, Frequency x Duration interaction | none | Exp2a-c-PSID-Acc-FreqXDuration.jasp | | (Excluded 1 subject: 5304786 from 1080Bias) |
| **3a-b** | Across Exp 3a and Exp 3b, we find an equal and opposite effect of frequency probability, with the more-probable tone detected on average ~6% more accurately than the low probability tone (**Fig 5**; Frequency x Probability interaction, $p = 3.361 \times 10^{-6}$). | Repeated Measures ANOVA, Frequency x Experiment interaction | none | Exp3a-b-TPS-AccSummary240508.jasp | | |
| **4a-b** | In Exp 4a and Exp 4b, RTs to the high probability tone frequency are ~28 ms faster, on average, than those to the low-probability frequency ($p = 1.375 \times 10^{-6}$). | Repeated Measures ANOVA, Frequency x FreqProbability interaction | none | Exp4a-b-PSID-RT-FreqXDur.jasp | | |
| **4a-b** | We also observe the perceptual bias of Exp 2 even in this impoverished probability distribution, with faster RTs for long-low/short-high duration-to-frequency pairings (**Fig S1**; Frequency x Duration interaction, RT: $p = 9.34 \times 10^{-6}$; | Repeated Measures ANOVA, Frequency x Duration interaction | none | Exp4a-b-PSID-RT-FreqXDur.jasp | | |
| **4a-b** | Acc: $p = 6.318 \times 10^{-5}$ | Repeated Measures ANOVA, Frequency x Duration interaction | none | Exp4a-b-PSID-Acc-FreqXDur.jasp | | |

| | | | | | | |
|---|---|---|---|---|---|---|
| **3c** | In the statistically volatile context established by Exp 3c, there is a detection advantage for the more probable frequency, with significant 'flips' in detection accuracy related to short-term reversals in tone probability for the first three blocks of Exp 3c (Fig 5; Frequency x Block interaction, p = 2.495 x 10-5, each block $p < 0.05$ ). In the final block, there is no significant difference in detection accuracy across frequencies (although numerically the trend is in the expected direction; all other blocks, $p < 0.05$). | Repeated Measures ANOVA, Frequency x TimeByQuarter | linear contrasts for each block comparison of the two conditions, bonferroni corrected | Exp3c-TPS-AccSummary240508.jasp | | |
| **4c** | Likewise, transient changes in probability distribution affect the efficiency of perceptual decisions in Exp 4c (Frequency x Block interaction, $p = 5.253 \times 10^{-7}$). RTs are fastest for the more probable frequency in Blocks 1, 3, and 4 (all $p < 0.04$ Bonferonni-corrected). | RM-ANOVA, Freq x Block x Duration; Freq x Block interaction | linear contrasts for each block comparison of the two conditions, bonferroni corrected | Exp4c-PSID-RT-FreqXDur.jasp | | (Excluded 3 subjects: 6259715, 6259743, 6259980) |
| **4c** | Even in this dynamic context we observe the systematic frequency-duration perceptual bias discovered in Exp 2 (**Fig S1**; Frequency x Duration interaction, RT: p = 0.019; Acc: p = 0.019). | RM ANOVA, Freq x Block x Duration (RT), interaction Frequency x Duration; Repeated Measures ANOVA, Frequency x Duration interaction, (Acc) | | Exp4c-PSID-Acc-FreqXDur.jasp, Exp4c-PSID-RT-FreqXDur.jasp | | (Excluded 3 subjects: 6259715, 6259743, 6259980) |
| **5a** | With equiprobable frequencies in the first half of Exp 5a (**Fig 2**), detection accuracy is relatively consistent across frequency (**Fig 6**; overall ~65%, with unexpectedly better detection for 800 Hz, p = 0.009). | Exp5a-FirstHalfOnlyEquiprobable-MakeSureToToggleExpt; FrequencyFirstHalfOnl | Exp5a-FirstHalfOnlyEquiprobable-MakeSureToToggleExpt | Exp5a-b-ByFreq.jasp | Filter to be only 5a (equi first) | |
| **5a** | In the second half of Exp 5a, probabilities shift to mirror Exp 1b (1000 Hz 75%; all others 6.25%). This probability shift drives differential changes in accuracy ($p = 8.511 \times 10^{-7}$). | Repeated Measures ANOVA, DistFromCentre x ExptHalf interaction | | Exp5a-TPS-EquiFirst-Acc-SummaryByDist.jasp | | |

| | | | | |
|---|---|---|---|---|
| **5a** | The now-higher-probability 1000 Hz tones are better detected than they were in the first (equiprobable) half of Exp 5a (p = 0.013, whereas the now-less-probable frequencies nearest (p = 0.041) and furthest (p = 0.004) from 1000 Hz are more poorly detected. | Repeated Measures ANOVA, Linear contrasts, Bonferroni-corrected | | Exp5a-TPS-EquiFirst-Acc-SummaryByDist.jasp |
| **5b** | In Exp 5b, we reverse distribution order (**Fig 2**). With a unimodal distribution centered on 1000 Hz in the first half of Exp 5b, detection generally resembles Exp 1c (**Fig 6**), with better accuracy for high-probability 1000 Hz compared to low-probability frequencies ($p = 2.77 \times 10^{-10}$), but with only a numerical detection advantage for frequencies nearest (920 and 1080 Hz) versus furthest (800 and 1200 Hz) from the probable center frequency (p = 0.312, Bonferroni-corrected). | RM-ANOVA,UnimodalFirstHalfOnly | first contrast is high versu low probability, second contrast is near versus far, Bonf corrected | Exp5b-TPS-80-20First-Acc-SummaryByDist.jasp |
| **5b** | When tone frequencies become equiprobable mid-study, again the probability shift drives differential changes in accuracy ($p = 1.815 \times 10^{-4}$). | Repeated Measures ANOVA | DistFromCentreX ExptHalf interaction | Exp5b-TPS-80-20First-Acc-SummaryByDist.jasp |
| **5b** | Here, the influence of the unimodal distribution carries over to confer a detection advantage to the formerly-highly-probable 1000 Hz compared to formerly low-probable-frequencies ($p = 1.068 \times 10^{-5}$), | RM-ANOVA,Equiprobable 2ndHalfOnly | first contrast (center vs near/far), Bonf corrected | Exp5b-TPS-80-20First-Acc-SummaryByDist.jasp |
| **5b** | although detection accuracy of 1000 Hz tones decreases from the first to the second study half (p = 0.0035). | Repeated Measures ANOVA | 3rd contrast (Centre First Minus Centre Second), Bonf corrected | Exp5b-TPS-80-20First-Acc-SummaryByDist.jasp |
| **5b** | There is not a commensurate improvement of detection for formerly low-probability tones, despite a more than 3-fold probability increase (p = 1, Bonferroni corrected). | Repeated Measures ANOVA | 5th contrast, Bonf corrected | Exp5b-TPS-80-20First-Acc-SummaryByDist.jasp |

| | | | | | | |
|---|---|---|---|---|---|---|
| **6a** | In the first half of Exp 6a, duration decision RTs across equiprobable frequencies are similar (**Fig 6**, p = 0.163). | RM-ANOVA-6aONLYFirstHalfOnly-Toggle! | Exclude BiasFirst; main effect of frequency | Exp6a-b-RT-FreqXExptHalfXDurationFINAL.jasp | | Excluded 2 subjects: 6516557, 6517370) |
| **6a** | When probabilities shift to a unimodal distribution centered on 1000 Hz at mid-study, RTs drop overall (p = 0.011) | Exp6a-ONLY-MakeSureToToggle!; | Exclude BiasFirst; main effect of Expt Half | Exp6a-b-PSID-RT-DistXExptHalf.jasp | | Excluded 2 subjects: 6516557, 6517370) |
| **6a** | Although there is a numerical RT advantage for the now-probable 1000 Hz compared to more distant frequencies, this pattern does not differ significantly across experiment half (p = 0.245). | Exp6a-ONLY-MakeSureToToggle!; | Exclude BiasFirst; interaction of Expt Half and DistanceFromCenter | Exp6a-b-PSID-RT-DistXExptHalf.jasp | | Excluded 2 subjects: 6516557, 6517370) |
| **6b** | In the first, unimodal probability half of Exp 6b, duration decisions exhibit the "V" shape around the high-probability 1000 Hz tone also observed in Exp 2b (effect of frequency, p =6.847 x 10-8, Fig 6). | RM-ANOVA-6aONLY-Toggle! | Exclude EquiFirst, main effect of frequency | Exp6a-b-RT-FreqXExptHalfXDurationFINAL.jasp | | Excluded 1 subject: 7100964) |
| **6b** | Decisions about low-probability frequencies near to 1000 Hz are slowed compared to 1000 Hz itself (p = 0.024) but faster than to those further away from 1000 Hz (p = 0.004). | Exp6b-ONLYFirstHalfOnly-MakeSureToToggle! | ExcludeEquiFirst, contrasts 1 & 2 (FreqDistance)bonf corrected for 2 tests | Exp6a-b-PSID-RT-DistXExptHalf.jasp | | Excluded 1 subject: 7100964) |
| **6b** | When all frequencies become equiprobable mid-study in Exp 6b, there is a change in the degree to which frequency modulates duration decisions (p = 0.024), | RM-ANOVA-Expt6bOnly-TOGGLE! | ExcludeEquiFirst, Frequency x ExptHalf interaction | Exp6a-b-RT-FreqXExptHalfXDurationFINAL.jasp | | Excluded 1 subject: 7100964) |
| **6b** | Even though 1000 Hz is now 20% probable, RTs are not significantly different than in the first experiment half when it was 80% probable (p = 0.796). | Exp6b-ONLY-MakeSureToToggle! | ExcludeEquiFirst, Contrast 1 | Exp6a-b-PSID-RT-DistXExptHalf.jasp | | Excluded 1 subject: 7100964) |
| **6b** | Like detection in Exp 5b, there is carryover from experience with the unimodal distribution in the first half of the study, such that duration decision RTs are still modulated by frequency (p = 8.306 x 10^-5). | Exp6b-ONLY2ndHalfOnly-MakeSureToToggle! | ExcludeEquiFirst, Frequency main effect | Exp6a-b-RT-FreqXExptHalfXDurationFINAL.jasp | | Excluded 1 subject: 7100964) |

| | | | | | | |
|---|---|---|---|---|---|---|
| **6b** | RTs to report decisions for 1000 Hz continue to be significantly faster than for the now-equally-probable far frequencies ($p = 0.003$), although not significantly faster than nearby frequencies ($p = 0.405$). | Exp6b-ONLY2ndHalfOnly-MakeSureToToggle! | ExcludeEquiFirst, Contrasts 2 and 1, bonferroni corrected | Exp6a-b-PSID-RT-DistXExptHalf.jasp | | Excluded 1 subject: 7100964) |
| **6a-b** | Finally, we again observe the duration-frequency bias established in the prior duration decision studies (**Fig S1**, Frequency x Duration interaction, RT: $p = 1.608 \times 10^{-4}$; | Repeated Measures ANOVA | Interaction of Frequency and Duration | Exp6a-b-RT-FreqXExptHalfXDurationFINAL.jasp | | Excluded 3 subjects: See above) |
| **6a-b** | Finally, we again observe the duration-frequency bias established in the prior duration decision studies (Fig S1, Frequency x Duration interaction … Acc: $p = 0.006$). | Repeated Measures ANOVA | Interaction of Frequency and Duration | Exp6a-b-Acc-FreqXExptHalfXDuration.jasp | | Excluded 3 subjects: See above) |
| **7a-c-e** | In Exp 7a,c,e, an equiprobable distribution precedes a switch to a unimodal distribution centered on 1000 Hz. Across these three studies, detection accuracy in the equiprobable first halves does not vary across frequency ($p = 0.393$) nor is it modulated by the different frequency ranges across Exp 7a,c,e ($p = 0.115$) and there is no interaction of frequency and range ($p = 0.119$). | Range Expts, EquiFirstOnly, FirstHalfOnly | Main effect of DistFromCenter, Main effect of Range, interaction of frequency and range; exclude all Bias conditions | Exp7a-c-e-EquifirstXDist.jasp | | |
| **7a-c-e + 5a** | Average detection accuracy across these equiprobable distributions is 64%, which does not differ from that in the 5-frequency distribution of Exp 5 ($p = 0.219$). | ANOVA (note, these data are averaged across all frequences) | Main effect of Experiment | Exp7a-c-e-PlusExp5a-FirstHalfOnlyAvgFreq.jasp | | |
| **7a-c-e** | The introduction of the unimodal distribution differentially effects frequency detection, depending on distance from 1000 Hz ($p = 1.622 \times 10^{-11}$). | RM ANOVA on Range Expts, Equiprobable first only | Interaction of ExptHalf and DistFromCenter | Exp7a-c-e-EquifirstXDist.jasp | | |

| | | | | |
|---|---|---|---|---|
| **7a-c-e** | When 1000 Hz shifts from equiprobable (14.3%) to highly probable (71.4%), there is a small but reliable *increase* in detection accuracy (p = 0.002). It is notable that this five-fold increase in probability (and ~16-fold increase in relative probability compared to low-probability frequencies) only confers an average 3.7% detection improvement. This mild enhancement is not significantly influenced by the range of frequencies (p = 0.365). | Range Expts, EquiFirst Only, CenterFreq Only | Main effect of Expt Half, and Interaction of Range x ExptHalf | Exp7a-c-e-EquifirstXDist.jasp |
| **7a-c-e** | Examining the off-center frequencies that drop in probability (14.3% to 4.8%) upon introduction of a unimodal distribution, we observe a significant *decrease* in detection accuracy of 4.7% ($p = 4.798 \times 10^{-9}$), the magnitude of which does not differ significantly across range (p = 0.337). | Range Expts, EquiFirstOnly, ExcludeCenterFreq | Main effect of Expt Half, and Interaction of Range x ExptHalf | Exp7a-c-e-EquifirstXDist.jasp |
| **7b-d-f** | Turning next to Exp 7b,d,f (**Fig 2x**), what happens when initial experience with a unimodal distribution shifts mid-study to equiprobable presentation? As now expected from prior results, detection of the high-probability unimodal center frequency is considerably more accurate than detection of improbable frequencies ($p = 1.220 \times 10^{-40}$). | RangeExpts,Unimodal(Bias)First,FirstHalfOnly | Contrast - FirstHalfOnlyFreq | Exp7b-d-f-RangeExpts.jasp |
| **7b-d-f** | Detection of low-probability frequencies is modulated by proximity to the high-probability center frequency (p = 0.010); accuracy is higher for frequencies nearest the high-probability center frequency compared those at middle (p = 0.023) or far frequencies (p = 0.023). | RangeExpts,Unimodal(Bias)First,FirstHalfOnly,NoCenter | Main effect of 'FirstHalfOnlyFreqNoCenter'; Post Hoc comparisons, holm-correct p-values, Int vs Near, Far vs Near [Note, post hoc table order is confusing!] | Exp7b-d-f-RangeExpts.jasp |

| | | | | |
|---|---|---|---|---|
| **7b-d-f** | However, the relatively preserved detection accuracy for tones near the high-probability frequency compared to those is observed only in Exp 7b for the narrow range (near vs. middle, p = 0.017, near vs. far, p = 4.449 x $10^{-4}$). | RangeExpts,Unimodal (Bias)First,FirstHalfOnly,NoCenter | Contrasts 1-4, contrast 1 is Narrow range, Near - Int; contrast 4 is Narrow range, Near - far. Values are bonferroni corrected for the 4 comparisons | Exp7b-d-f-RangeExpts.jasp |
| **7b-d-f** | The effects on detection of proximity to the high-probability 1000 Hz are modulated by the switch to an equiprobable distribution (p = 3.279 x $10^{-11}$). | RM ANOVA on Range Expts, Unimodal (Bias) First OMNIBUS | Interaction of ExptHalf and DistFromCenter | Exp7b-d-f-RangeExpts.jasp |
| **7b-d-f** | We observe a continued, but smaller, detection advantage for the formerly high-probability center frequency compared to formerly improbable frequencies (1.066 x $10^{-14}$). | RangeExpts,Unimodal (Bias)First,SecondHalfOnly | Contrast 1 | Exp7b-d-f-RangeExpts.jasp |
| **7b-d-f** | This change is driven by a *decrease* (difference of 7.1, p = 1.137 x $10^{-12}$) in detection accuracy for the center frequency as it becomes 5 times less probable, as well as a smaller (difference of ~2%, p = 0.007) *increase* in accuracy as off-center frequencies become 3 times more probable (**Fig 2x**). | RM-ANOVA, RangeExpts, Bias-First. 1kHz only, AND Range Expts, Unimodal (Bias) First OMNIBUS, NoCentreFreq | Main effect of Expt Half for both | Exp7b-d-f-RangeExpts.jasp |
| **7b-d-f** | This residual advantage does not vary significantly with distance from the center frequency (p = 0.213) or interact with the range of frequencies presented (p = 0.202). | RM-ANOVA, RangeExpts, Bias-First. NoCentreFreq | Interaction of ExptHalf x DistFromCenter, ExptHalf x DistFromCenter x Range | Exp7b-d-f-RangeExpts.jasp |

| | | | | |
|---|---|---|---|---|
| **7a-f, unimodal halves only** | Next, we ask if hysteresis is also observed in detection accuracy for 1000 Hz in a unimodal distribution *after* prolonged initial exposure to an equiprobable distribution (second half of Exp 7a,c,e) compared to when the experiment begins with a unimodal distribution (first half of Exp 7b,d,f). We find that pre-exposure to 336 trials of the flat probability distribution diminishes detection rates for the high-probability 1000 Hz tone in the subsequent unimodal distribution by 5.8% relative to when the identical unimodal distribution is encountered first ($p = 6.394 \times 10^{-4}$). The persistent damping effect of encountering the equiprobable distribution is not significantly affected by the range of frequencies encountered ($p = 0.768$). | 'RangeExpts-UnimodaHalvesCombined-CentreFreqOnly' | Main effect of Order | Exp7a-f-RangeExpts-BiasedHalvesOnlyXDistFromTargAcc.jasp |
| **7a-f, unimodal halves only** | Finally, we aggregate detection data for off-center frequencies across the unimodal conditions from Exp 7a,c,e (for which it was second) and Exp 7b,d,f (for which it was first) to maximize the power to detect influences of frequency range and distance from the higher-probability center frequency. Frequency range has an effect on detection in unimodal probability distributions ($p = 0.005$). Specifically, a wide frequency range impairs overall off-center detection accuracy, compared to when the frequency range is narrow ($p = 0.006$). (The middle frequency range falls in-between and differs significantly from detection in wide, $p = 0.037$, but not narrow, $p = 0.429$, ranges.) | RangeExpts-UnimodalHalvesCombined-NoCentreFreq | Main effect of Range, condition-wise comparisons are Holm-corrected post hoc tests | Exp7a-f-RangeExpts-BiasedHalvesOnlyXDistFromTargAcc.jasp |

| | | | | |
|---|---|---|---|---|
| **7a-f, unimodal halves only** | Moreover, the shape of the drop-off in detection accuracy from the high-probability center frequency is significantly graded only in the narrow frequency range, between near and mid frequency bands condition (p = 0.013), with a non-significant decrease between middle and far frequencies (p = 0.318). | RangeExpts-UnimodalHalvesCombined-NoCentreFreq | Contrasts 1-4, contrast 4 is near vs middle (narrow), contrast 3 is middle vs far (narrow); p-values Bonferroni-corrected for 4 comparisons | Exp7a-f-RangeExpts-BiasedHalvesOnlyXDistFromTargAcc.jasp |
| **8b** | We first ask whether the consistent experience with 1000 Hz in the first half of Exp 8b yields accumulating detection accuracy improvements (**Fig 8**). It does not: accuracy in the first quarter of trials is 78% (aligned with expectations from listener-specific thresholding) then decreases slightly to plateau at 75% for the remaining trials (p = 0.015). | CenterFreqOnlyFirstHalfByBlock | Main effect of block pair | Exp8b-21Freq1kHzFirstSubsByDistAndBlockPair.jasp |
| **8a, 1a** | Similarly, neither Exp 1a (p = 0.210) or Exp 8a (p = 0.451) exhibit improved detection across a block of trials with only 1000 Hz tones. | Repeated Measures ANOVA (1a), CenterFreqByBlock2ndHalf (8a) | Main effect of block; main effect of block pair 2nd half | Exp1a-TPS-1kHz-ByBlock.jasp; Exp8a-21FreqEquiFirstSubsByDistAndBlockPair.jasp |
| **8a** | There is a similar initial detection decrement of ~5% across the first quarter of the 20-equiprobable-frequency trials of Exp 8a with no further change ($p = 9.669 \times 10^{-6}$). | OffCenterFreqsOnlyFirstHalfByBlockPair | Main effect, FirstHalfBlockPair | Exp8a-21FreqEquiFirstSubsByDistAndBlockPair.jasp |
| **7a,c,e** | This same pattern emerges in the initial equiprobable blocks of Exp 7a,c,e ($p = 1.375 \times 10^{-5}$). | Repeated Measures ANOVA | Main effect of Block | Exp7a-c-e-7FreqEquiFirstBlocksOnlyByBlock.jasp |
| **5a, 7a,c,e, 8a** | Detection accuracy for equiprobable distributions that are experienced in the first half of a study does not differ over experiments (Exp 5a, 7a,c,e, and 8a; p = 0.387). | ANOVA | Main effect of Experiment | Exp5a+Exp7a-c-e+Exp8a-7Freq+21FreqEquiFirstBlocksOnlyBySub.jasp |

| | | | | |
|---|---|---|---|---|
| **8a, 1a, 8b** | Turning next to the nature of the hypothesized attentional filter, we first examine whether initial experience with the 20-tone equiprobable distribution in Exp 8a (which does not include 1000 Hz) impacts subsequent detection in the 1000-Hz-only block. It does not: detection of 1000 Hz in Exp 8a did not differ from either Exp 1a ($p = 0.315$) or the first half of Exp 8b ($p = 0.837$), each of which involved blocks of trials with only 1000 Hz at the beginning of the study. | ANOVA | Tukey-corrected posthocs across experiment pairs | Exp1a+Exp8a-b1000HzOnlyPlus21Freq1000HzMeanAccBySub.jasp |
| **8a,b** | In contrast, massed exposure to 1000 Hz in the first half of Exp 8b drives a dramatic, long-lasting, and frequency-specific detection decrement for the subsequently encountered 20 equiprobable frequencies, as compared to detection across equiprobable frequencies in Exp 8a (interaction of Distance-from-1000-Hz x Exp, $p = 2.618 \times 10^{-4}$). | RM ANOVA, 20-freq Dist x order, no center frequency | Interaction, DistFromCenter x experiment | Exp8a-b-TPS-21Freq-AllSubsByDistFromCenter.jasp |
| **8a,b** | Specifically, as shown in **Fig 8**, detection of frequencies at far (2 to 3.9 semitones) and intermediate (1 to 2 semitones) distances from 1000 Hz were detected much less accurately after massed experienced with 1000 Hz (Exp 8b; far: $p = 1.668 \times 10^{-3}$, intermediate: $p = 9.007 \times 10^{-4}$), compared to equiprobable presentation at the beginning of the study (Exp 8a). This suppressive effect was rescued by proximity to the now-absent 1000 Hz in the second half of Exp 8b, with frequencies within about a semitone from 1000 Hz eliciting detection accuracies roughly on par with those from Exp 8a ($p = 0.362$). | RM ANOVA, 20-freq Dist x order, no center frequency | Contrasts 2, 3 & 4, bonferroni corrected for 4 comparisons (note order in table is different than text) | Exp8a-b-TPS-21Freq-AllSubsByDistFromCenter.jasp |

| | | | | |
|---|---|---|---|---|
| **8a,b** | One might expect that any initial learning across the 1000-Hz-only distribution would be overwhelmed by the mid-study shift to the high-uncertainty 20-frequency equiprobable distribution. However, we see the opposite: across the second half of Exp 8b, there is no significant change in overall detection accuracy (p = 0.165), nor any change across time in relative accuracy of detection across frequencies (**Fig 8**; p = 0.568). The large advantage for detection of frequencies near 1000 Hz compared to intermediate and far frequencies persists to the final 80 trials of Exp 8b (p = 0.006, **Fig 8**). | NonCenterFreqsByBlockIn2ndHalf | Main effect of BlockPair2ndHalf; Interaction BlockPair2ndHalf x DistFromCenter; contrast 1 (last block pair) | Exp8b-21Freq1kHzFirstSubsByDistAndBlockPair.jasp |
| **8a,b** | Here, there is strong suppression of frequencies far and intermediate distances from 1000 Hz in Exp 8b compared to detection of the same frequencies in the equiprobable half of Exp 8a. As with the within-experiment comparison, this difference is observed through the entirety of the second half of the study, again extending even to the last quarter of trials (p = 0.009). | Repeated Measures ANOVA | Contrast 1, comparison of 2 conditions in last block pair | Exp8a-b-21Freq-Cross-ExptCompareFarIntOverBlockPairs.jasp |

#### Fig. S1. A novel frequency-duration perceptual bias

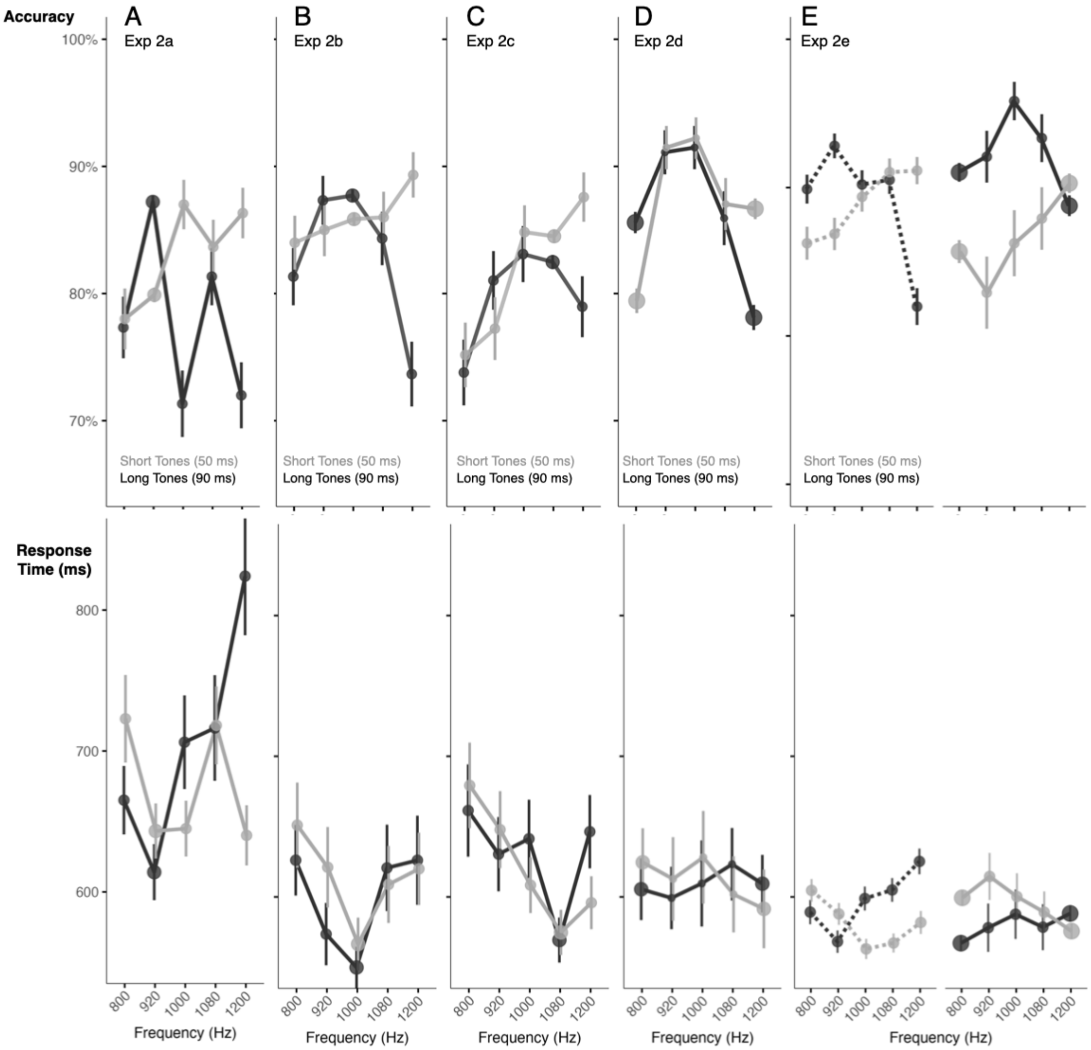

**Figure S1. A novel frequency-duration perceptual bias.** Each panel plots mean duration decision accuracy as a function of acoustic frequency for Exp 2a-c, respectively (Panels A, B, C). Error bars reflect standard error of the mean. Responses to short (50 ms) tones are plotted in grey and responses to long (90 ms) tones are plotted in black. Marker size indicates tone probability. Duration decisions for lower-frequency tones (800, 920 Hz) are more accurate and faster for long (90 ms) compared to short (50 ms) whereas those for the highest tone frequency (1200 Hz) are more accurate and faster for short compared to long tones. This underlying perceptual bias is greatest at the edges of the distribution range, interacting substantially with the bimodal distributions of Exp 2d-e. This helps to explain the absence of a dual spotlight influence of bimodal distributions in Exp 2d-e that mirrors those observed for detection in Exp 1e-f.Fig. S2. Type or paste legend here. Paste figure above the legend.

**Fig. S2. The influence of experience with a single frequency (point distribution) persists across the second half of the study**

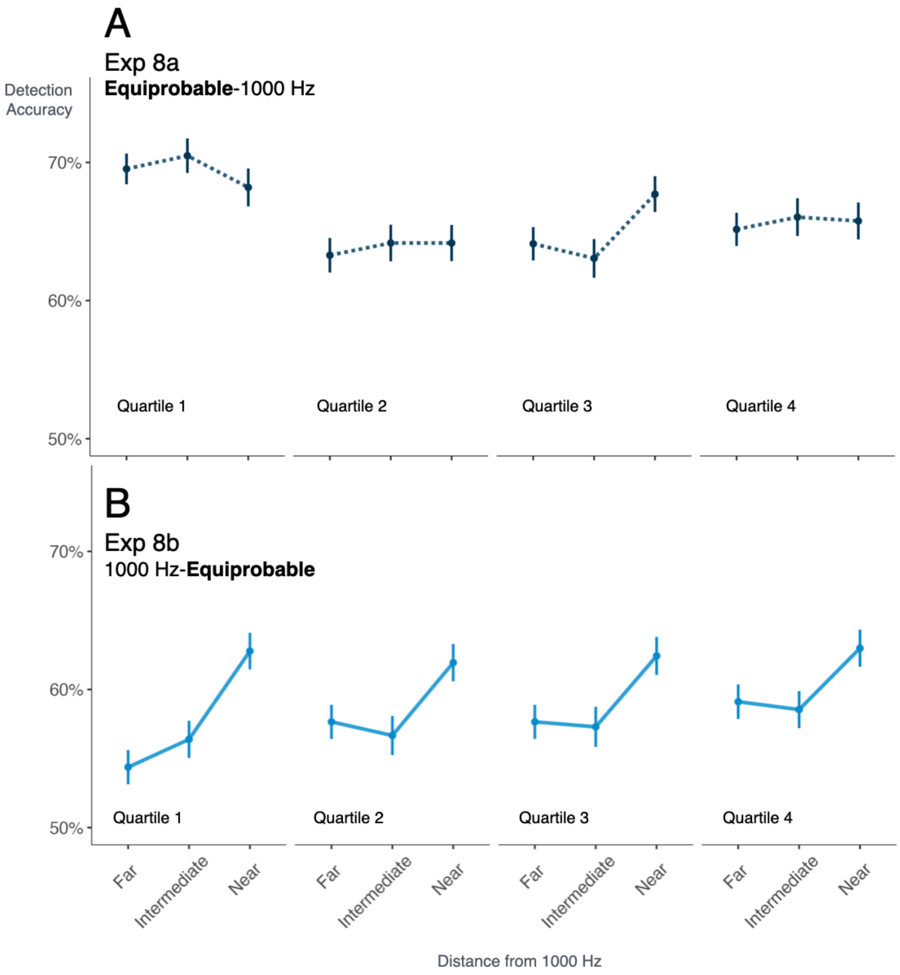

**Fig S2. The influence of experience with a single frequency (point distribution) persists across the second half of the study.** Each panel plots mean detection accuracy across equiprobable distributions (first half of experiment in Exp 8a, second half of experiment in Exp 8b) as a function of distance of tone frequency from 1000 Hz (near, intermediate, far) and quartile of trials within the half of the experiment involving the equiprobable distribution. Error bars correspond to standard error of the mean. **(A)** Exp 8a provides an estimate of baseline detection accuracy across the 20 frequencies in the first half of the study. (B) In contrast, detection accuracy in Exp 8b — which follows experience with 1000 Hz only in the first half of the study — is significantly suppressed for tones with frequencies intermediate and far distances from 1000 Hz. Note that this persists even into the final quartile (80 trials) of Exp 8b. The absence of 1000 Hz in these equiprobable distributions rules out trial-wise perceptual interactions or a relative probability difference for the lingering influence of the point distribution experienced at 1000 Hz in the first half of Exp 8b.
